# A genome-wide knock-out screen for actors of epigenetic silencing reveals new regulators of germline genes and 2-cell like cell state

**DOI:** 10.1101/2021.05.03.442415

**Authors:** Nikhil Gupta, Lounis Yakhou, Julien Richard Albert, Fumihito Miura, Laure Ferry, Olivier Kirsh, Marthe Laisné, Kosuke Yamaguchi, Cécilia Domrane, Frédéric Bonhomme, Arpita Sarkar, Marine Delagrange, Bertrand Ducos, Maxim V. C. Greenberg, Gael Cristofari, Sebastian Bultmann, Takashi Ito, Pierre-Antoine Defossez

## Abstract

Epigenetic mechanisms are essential to establish and safeguard cellular identities in mammals. They dynamically regulate the expression of genes, transposable elements, and higher-order chromatin structures. Expectedly, these chromatin marks are indispensable for mammalian development and alterations often lead to diseases such as cancer. Molecularly, epigenetic mechanisms rely on factors to establish patterns, interpret them into a transcriptional output, and maintain them across cell divisions. A global picture of these phenomena has started to emerge over the years, yet many of the molecular actors remain to be discovered. In this context, we have developed a reporter system sensitive to epigenetic perturbations to report on repressive pathways based on *Dazl,* which is normally repressed in mouse ES cells. We used this system for a genome-wide CRISPR knock-out screen, which yielded expected hits (DNMT1, UHRF1, MGA), as well as novel candidates. We prioritized the candidates by secondary screens, and led further experiments on 6 of them: ZBTB14, KDM5C, SPOP, MCM3AP, BEND3, and KMT2D. Our results show that all 6 candidates regulate the expression of germline genes. In addition, we find that removal of ZBTB14, KDM5C, SPOP and MCM3AP led to similar transcriptional responses, including a reactivation of the 2-cell like cell (2CLC) signature. Therefore, our genetic screen has identified new regulators of key cellular states.

## Introduction

Embryonic development starts with a single cell, the zygote. By nature, the zygote is totipotent: it can give rise to all cells of the embryo. This totipotency is still present in the 2-cell embryo, as each cell can regenerate a whole embryo and its extra-embryonic tissues, but then it quickly wanes with ensuing divisions. By the morula stage, cells of the inner cell mass are no longer totipotent, but only pluripotent: they can generate any embryonic cell lineage, including the germline, but cannot contribute to extra-embryonic tissues (Beddington & Robertson, 1989). Cells with different potencies acquire a unique cell identity. Different epigenetic pathways are key contributors to the regulation of gene expression and the maintenance of cellular identity. Some of the main molecular actors involved in shaping the epigenetic landscape include DNA methylation, histone modifications, and chromatin proteins such as the Polycomb repressive complex (PRC) (Schübeler et al., 2015; Blackledge et al., 2015). A complex interplay is at work across these chromatin modifications finely tuned over time, and dynamically responds to the multiple environmental cues received by the cell. This is of paramount importance in mammalian development and disease, as disturbing these patterns often leads to embryonic lethality or tumorigenesis (Greenberg & Bourc’his, 2019; Piunti & Shilatifard, 2020).

DNA methylation is the covalent modification of DNA by a methyl group (CH_3_) at the C5 position, yielding 5-methylcytosine mostly in the context of CpG dinucleotides (Li & Zhang, 2014). The establishment and maintenance of genomic DNA methylation patterns are carried out by the DNA methyltransferase (DNMT) family and accessory proteins such as UHRF1 (Ferry et al., 2017; Laisné et al., 2018; Petryk et al., 2020), creating a pattern that is stable within a cell type and presents reproducible differences between varying cell types (Schultz et al., 2015). DNA methylation is essential in mammals: genetic ablation of DNMT1 or UHRF1 results in a progressive loss of DNA methylation and embryonic lethality in mammals (Li et al., 1992). DNA methylation at promoters, gene bodies, and many transposable elements has a causal role in the regulation of gene expression (Schübeler et al., 2015). Other covalent epigenetic modifications include histone post-translational modifications (PTMs) such as methylation, acetylation, ubiquitination, and are associated with gene repression or activation. They are deposited or removed by specific writers/erasers to create a chromatin context able to recruit or repel transcription factors at the promoters and enhancers (Morgan & Shilatifard, 2020).

A specific class of genes, germline genes which include testis-specific genes, have been demonstrated to be regulated by both DNA methylation and PRC. However, the extent of regulation differs based on cell origin (Borgel et al., 2010; Hargan-Calvopina et al., 2016; Maezawa et al., 2017). Repression of germline genes outside of their developmental window is important for the maintenance of cell identity and often misexpressed in several cancers and syndromes (Gibbs & Whitehurst, 2018). Remarkably, in primordial germ cells, the germline genes involved in genome defense against transposable elements are mainly regulated by DNA methylation (Hackett et al., 2012), and are activated when DNA methylation reprogramming occurs during development (Greenberg & Bourc’his, 2019). However, other repressive pathways repress germline genes, both in somatic tissues and embryonic stem cells, either directly or synergistically with the DNA methylation machinery (Endoh et al., 2017; Hackett et al., 2012; Hill et al., 2018). Of note, members of the ncPRC1.6 complex (E2F6, MAX) can recruit DNMTs to synergistically repress *Dazl* and *Taf7l* loci (Velasco et al., 2010; Tatsumi et al., 2018) while other genes (*1700013H16Rik*, *Asz1*, *Sycp1*) are repressed by DNA methylation only (Hill et al., 2018).

Nevertheless, the picture is still far from complete, and discoveries continue to be made that clarify and/or explain the process. Factors contributing to the deposition, the maintenance, the erasure, or the interpretation of epigenetic marks are still incompletely known. To obtain a full molecular understanding of the process, we performed a CRISPR-based genetics screen with a reporter system in mouse embryonic stem cells (ESCs). Mouse ESCs are self-renewing pluripotent cell lines derived from the inner cell mass of the mouse preimplantation embryo (Evans & Kaufman, 1981), and are maintained in an undifferentiated state by the addition of the cytokine leukemia inhibitory factor (LIF) to the culture medium (Ying et al., 2008). Epigenetic state in mouse ESCs depends on culture conditions, cells grown in 2i/VitaminC/LIF medium (hereafter referred to as “2i”) have lower DNA methylation than cells grown in serum/LIF medium (hereafter referred to as “serum”) (Ficz et al., 2013; Blaschke et al., 2013; Habibi et al., 2013) and undergo a reconfiguration of their repressive chromatin landscape (van Mierlo et al., 2019).

The mouse ESCs, like their counterparts in the embryo, are pluripotent but not totipotent. However, at a very low frequency, totipotent-like cells do arise in ES culture, where they can constitute ∼0.1-0.5% of the population (Macfarlan et al., 2012; Morgani et al., 2013). They share properties and markers with the 2-cell embryo and are therefore called “2-cell-like-cells” or 2CLCs. Building on this pivotal observation, several 2CLC markers have been discovered (Genet & Torres-Padilla, 2020): 2CLCs are characterized by the derepression of the MERVL retrotransposons, which in part drive the 2-cell transcriptional program. This program contains genes such as the family of Zinc finger and SCAN domain proteins *Zscan4a-f* (Falco et al., 2007) and *Zfp352*. A master regulator of the 2CLC transcriptome is the transcription factor DUX (*Dux* in mouse/DUX4 in human), which binds and activates MERVL repeats and many promoters in this program (De Iaco et al., 2017; Hendrickson et al., 2017; Whiddon et al., 2017).

In this study, we focused our attention on repressive factors by selecting cells activating our epigenetic-sensor reporter system after gene knock-out, while recent genome-wide genetic studies screened for activating factors (Alda-Catalinas et al., 2020; Gretarsson & Hackett, 2020; Dixon et al., 2021). We successfully identified and validated novel epigenetic regulators, playing an essential role in the regulation of testis-specific genes, repeat elements, and cell state transition to 2CLC.

## Results

### Design and validation of the epigenetic reporter

Epigenetic silencing is dynamic in mouse ES cells and responds to culture conditions. Correspondingly, these changes activate a number of genes that are epigenetically repressed in serum condition but expressed in 2i (Figure 1A). The principle of our screen was to select one such gene and replace it with selectable markers, so as to carry out a CRISPR KO screen in serum condition, and to positively select the cells in which the KO caused a loss of silencing (Figure 1A). To select the gene in question, we considered 3 criteria: mechanism(s) of epigenetic repression, biological function(s) of the gene, and expression level in repressed and induced condition (to ensure high enough expression of the selection markers). These criteria led us to select *Dazl* as our target of choice. In serum condition, *Dazl* is repressed by two mechanisms: DNA methylation (Borgel et al., 2010; Hackett et al., 2012) and ncPRC1.6 (Endoh et al., 2017; Stielow et al., 2018); inactivation of either pathway causes its reactivation (Figure 1B). *Dazl* encodes an RNA-binding protein which is critical during gametogenesis (Zagore et al., 2018; Mikedis et al., 2020; Sharma et al., 2021), but it is also expressed during other developmental events, such as the 2-cell-stage (Rodriguez-Terrones et al., 2018) (Figure 1C). We verified by WGBS, RNA-seq, and RT-qPCR that the *Dazl* promoter was unmethylated and the gene expressed in 2i, whereas the *Dazl* promoter was methylated and the gene repressed in serum condition in our cellular background, J1 mouse ESC (Figure 1D-E).

**Figure 1:**
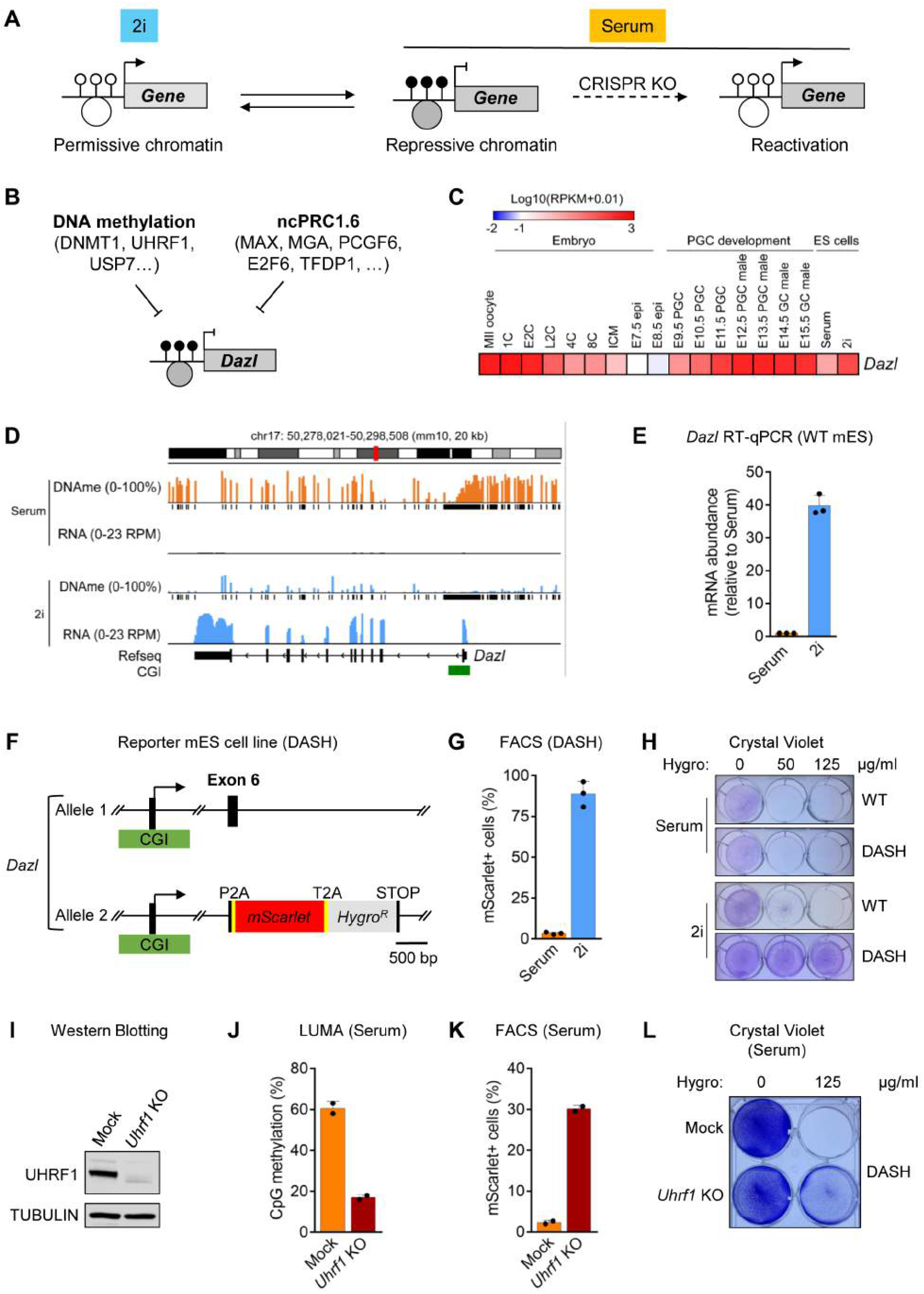
Screen design and generation of the DASH reporter cell line. (A) Principle of the screen: in ES cells, some genes are expressed in 2i condition (permissive chromatin and no DNA methylation at the promoter) but undergo transcriptional repression in serum condition (repressive chromatin and DNA methylation at the promoter). Knocking out factors essential for repression should triggers gene reactivation in serum condition. CpG DNA methylation status is represented by black (methylated) or white (unmethylated) lollipops. Chromatin status is depicted by grey (repressive) or white (permissive) circles. (B) *Dazl* is a suitable candidate, since, under serum condition, two synergistic inhibitory pathways contribute to *Dazl* repression: promoter DNA methylation and non-canonical PRC1.6 (ncPRC1.6). (C) Endogenous expression pattern of *Dazl* in early development. *Dazl* is a marker both of the 2-cell stage (E2C, L2C) and of primordial germ cell (PGC) development. (D) RNA-seq and WGBS in our cellular background confirm that *Dazl* is highly methylated and repressed in serum (non-permissive chromatin) and hypo-methylated and expressed in 2i (permissive chromatin). CGI: CpG island. (E) RT-qPCR showing the up-regulation of *Dazl* in mES cells cultured in 2i (relative to Serum condition). (F) The DASH mES reporter cell line. A reporter cassette is inserted into one of the *Dazl* alleles, it encodes two proteins separated by 2A self-cleaving peptides (P2A, T2A): the red-fluorescent mScarlet and the Hygromycin resistance enzyme (Hygro^R^). (G) DASH cells express mScarlet in 2i, but not in serum. FACS analysis (50,000 cells per condition in this figure and all following figures). (H) DASH cells are resistant to Hygromycin in 2i, but not in serum. (I) CRISPR KO of *Uhrf1* in DASH cells is confirmed by western blotting (J) LUMA confirms that *Uhrf1* KO in DASH cells leads to a global loss of DNA methylation. (K) *Uhrf1* KO in DASH cells results in mScarlet reactivation. (L) *Uhrf1* KO in DASH cells results in Hygromycin resistance; surviving cells stained with crystal violet.

We then used CRISPR/Cas9 to knock in 2 selectable markers in *Dazl*: mScarlet, a bright red fluorescent protein (Bindels et al., 2017), and Hygro^R^, which confers resistance to Hygromycin (Figures 1F, S1A). Hygro^R^ was chosen over other resistance markers because of its better tunability and compatibility with the expression level conferred by the *Dazl* promoter (Nakatake et al., 2013). The two markers were inserted in-frame in the *Dazl* coding sequence, at exon 6 (which is ∼6 kb away from the promoter and present in all splicing forms), and the two reporter proteins were separated by T2A and P2A self-cleaving peptides (Figures 1F, S1A). Recombinant clones were picked and characterized by genomic PCR and sequencing, one of the correct integrants was further analyzed, and Droplet digital PCR (ddPCR) was used to establish that only one mScarlet-Hygro^R^ insertion has occurred in the genome (Figure S1B). To summarize, we generated a heterozygous J1-based mES cell line in which one allele of *Dazl* is wild-type, and the other allele contains the mScarlet-Hygro^R^ insertion, which inactivates that allele (Figure 1F). We will refer to this line as the DASH (*Dazl*-Scarlet-Hygro) reporter line. As expected, DASH cells in 2i are mScarlet-positive (∼75% over threshold) and Hygromycin-resistant (to a dose > 125 µg/ml), but they are mScarlet-negative and Hygromycin-sensitive in serum condition (Figures 1G-H, S1C). Also as expected, the *Dazl* promoter was methylated in serum condition and demethylated in 2i (Figure S1D).

We next carried out a proof-of-concept experiment to verify that the selectable markers could be induced in serum upon the KO of an epigenetic repressor. For this, we used CRISPR to knock out a key factor in DNA methylation maintenance, UHRF1 (Figure 1I). As expected, the *Uhrf1* KO causes a global loss of DNA methylation as seen by LUMA (Figure 1J). The DASH cells grown in serum became mScarlet-positive and Hygromycin-resistant upon *Uhrf1* KO (Figures 1K-L), confirming the feasibility of a genetic screen.

### Genome-wide KO screening yields a list of 40 candidate hits

We grew the DASH cells in serum and infected them with a lentiviral library of ∼80,000 vectors co-expressing Cas9 and sgRNAs targeting ∼20,000 genes, the Brie library (Doench et al., 2016). The multiplicity of infection was ∼0.1, the coverage ∼150 infected cells per guide, and two independent screens were carried out in parallel. After an initial Puromycin selection to eliminate non-infected cells, Hygromycin selection was applied in two steps of increasing concentration (Figure 2A). mScarlet-positive cells represented ∼3% of the starting DASH population, this proportion increased to ∼25-30% of Hygromycin-resistant cells (Figure S2A), and these fluorescent cells were purified by FACS. By RT-qPCR we observed in this population a number of differences with the starting population: upregulation of the *Dazl* gene (as expected), but also increased expression of *Prdm14*, and decreased expression of *Dnmt3a* and *Dnmt3b*, all of which are markers of naïve cells (Figure S2B). The bulk DNA methylation in this selected population was lower than in the post-infection sample (Figure S2C), and the methylation of the *Dazl* promoter (assessed by MeDIP) was also lower than in the reference sample, indicating that some cells in the populations have lost DNA methylation locally and/or globally (Figure S2D). The sgRNAs present in these cells were amplified by PCR and sequenced by NGS, and the sequencing data statistically analyzed using MAGeCK (Li et al., 2014). This procedure yielded an ordered list of candidates, ranked by *p*-value (Figure 2B). The first 40 candidates had a *p*-value smaller than 5.10^−4^ and were analyzed further (Figure 2B). GO terms enriched in this set of candidates include “Maintenance of DNA methylation”, along with “Glycosaminoglycan synthesis” and “Heparan sulfate synthesis” (two related terms), and “FGFR signaling pathway” (Figure 2C). As expected, Uniprot GO terms “Repressor”, “Chromatin regulator”, and “DNA binding” were enriched (Figure S2E).

**Figure 2:**
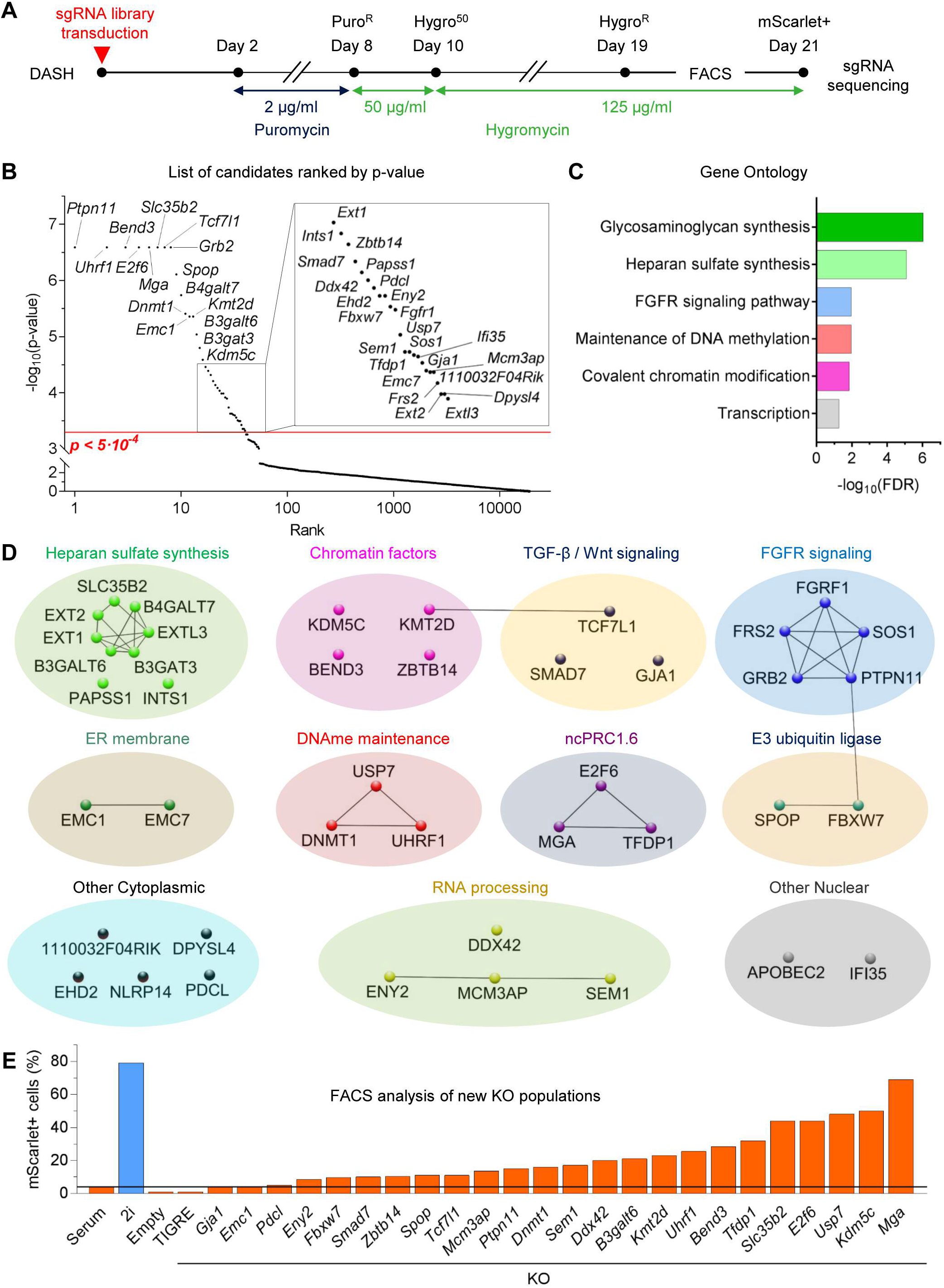
Genome-wide CRISPR knock-out screen for reporter gene reactivation in DASH mES cells. (A) Timeline of the screen. (B) Ranking of the hits by *p*-value. The top 40 hits (p < 0.0005) are above the red line. (C) Gene ontology (GO) terms (Biological Process) significantly enriched among the top 40 hits. (D) Protein-protein interaction network clustering of the top 40 hits generated using STRING analysis. (E) FACS analysis of mScarlet expression after KO of the indicated candidate genes. Cells transduced with an empty backbone, TIGRE KO and parental DASH cell lines (cultured in serum and 2i) served as controls.

We then grouped the 40 candidates on the basis of their known functions and their interactions in the STRING database (Figure 2D). We recovered 3 factors required for DNA methylation maintenance (UHRF1 ranked #2, DNMT1 ranked #11, USP7 ranked #28), as well as components of the ncPRC1.6 complex (E2F6 ranked #4, MGA ranked #5, TFDP1 ranked #38); these results validate the pertinence of our system as a reporter of epigenetic silencing. Besides these expected hits, we obtained 8 candidates in the “TGFß-Wnt signaling” and the “FGFR signaling” clusters; it is likely that these KOs genetically reproduce the effect of the 2i medium, rendering the cells more naïve, which could be the cause of *Dazl* reactivation. Similarly, mutation of genes in the “Heparan Sulfate/Glycosaminoglycan synthesis” has been reported to decrease FGF/MAPK signaling, and thus increase the proportion of naïve cells (Li et al., 2018). The top 40 hits also included genes involved in “RNA processing”, including all three members of the TREX-2 mRNA export complex (ENY2 ranked #24, SEM1 ranked #30, MCM3AP ranked #34), two histone-modifying enzymes (KMT2D ranked #13, KDM5C ranked #16), two E3-ubiquitin ligases (SPOP ranked #8, FBXW7 ranked #25) as well as additional nuclear factors (BEND3 ranked #3, ZBTB14 ranked #18).

We re-generated CRISPR KO populations for 24 selected genes in the top 40; of those 21 increased the number of mScarlet-positive cells in the population (Figure 2E). This 87.5% rate of true positives establishes the robustness of the screen results. As expected, we observed a large loss of DNA methylation at the *Dazl* promoter (Figure S2F) and throughout the genome (Figure S2G) after inactivation of UHRF1, DNMT1, and USP7 (Figure S2); smaller decreases were also observed in the other KO populations (Figures S2F-G).

### Screen validation rules out differentiation artifacts and global effects on known Dazl regulators

We then sought to prioritize the hits for further analyses. We expected to recover from the screen hits causing the cells to be blocked in differentiation, causing them to be “2i-like” even in serum condition, and therefore to have low DNA methylation and high *Dazl* expression. These hits can be identified on the basis of their expression of naïve markers, such as *Nanog* and *Prdm14*. In order to identify candidates in this class, we screened all 24 of our newly generated KO populations by Microfluidic RT-qPCR on a Fluidigm platform, examining the expression of 48 genes of interest (Figure 3A-C). Three hits showed elevated levels of *Nanog* and *Prdm14*, consistently with their known role in naïve pluripotency: PTPN11, SLC35B2, and B3GALT6 (Figure 3A). The *Dnmt1* KO showed the same expression pattern, again in line with previously published observations (Schmidt et al., 2012).

**Figure 3:**
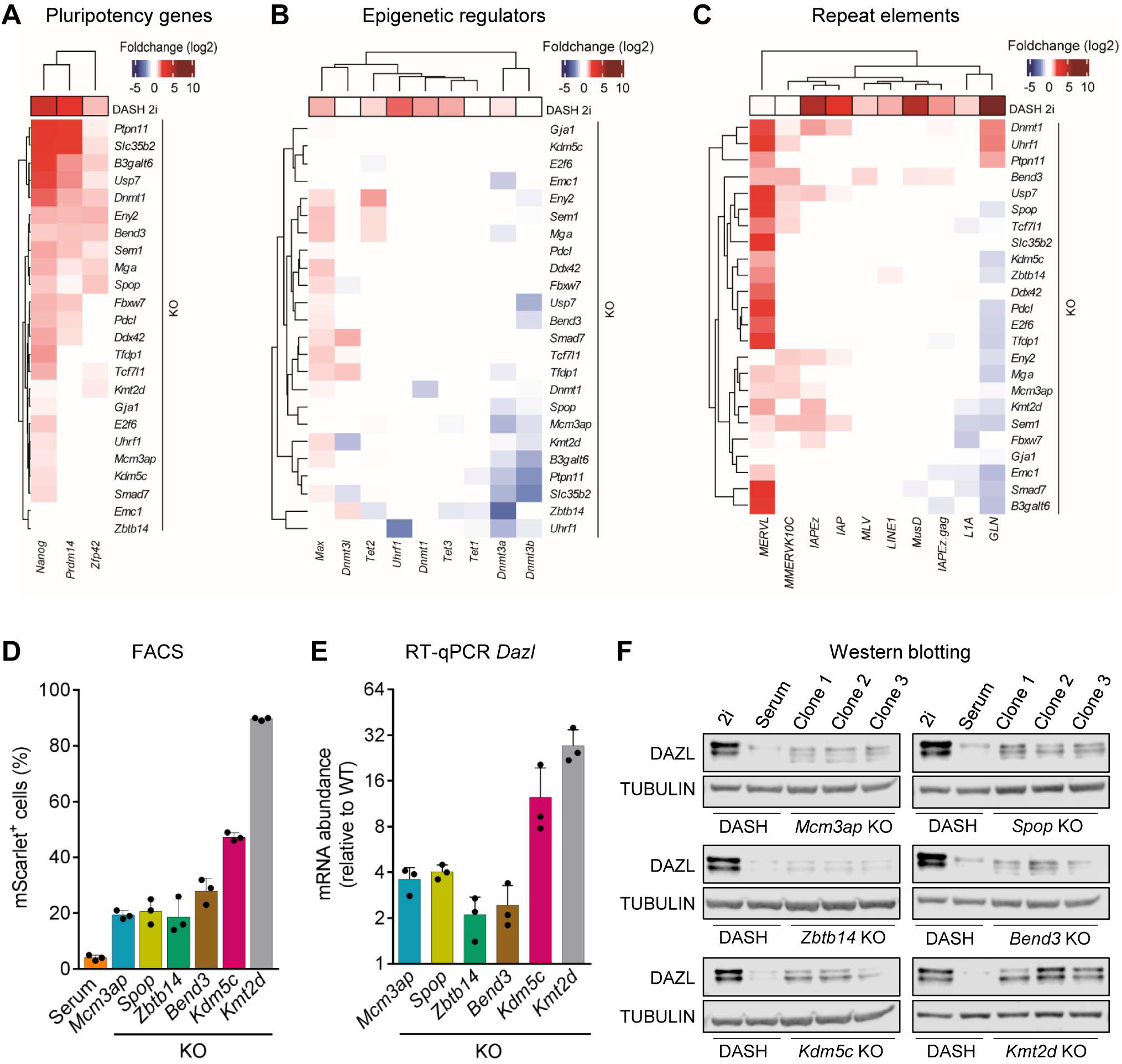
Secondary screens to characterize the regulators of Dazl. (A-C) DASH cells were infected with CRISPR vectors to KO the indicated genes (horizontal lines), selected for Hygromycin resistance, then used in Fluidigm assays on: (A) pluripotency genes, (B) epigenetic regulators, and (C) repeated elements. The average fold-change value of 2 technical replicates, relative to DASH mES cells cultured in Serum, is shown for each sample. (D) FACS analysis of the indicated KO populations, after Hygromycin selection. (E) RT-qPCR assay analysis of the indicated KO populations, after Hygromycin selection. (F) Western blot analysis of DAZL expression in the indicated cell populations. (D-F) Each data point represents an individual clonal cell line (n=3, clones per KO).

We then measured the expression of DNA methylation regulators (UHRF1, DNMT enzymes, DNMT3L, TET enzymes), in the different KOs (Figure 3B). Except for their corresponding KO samples, *Uhrf1* and *Dnmt1* levels were not changed, ruling out global repression of the methylation machinery as a likely mechanism for *Dazl* induction. Similarly, we found no major activation of *Tet1/2/3* (no global induction of the demethylation machinery) nor repression of *Max* in any of the KOs (Figure 3B). As expected, *Ptpn11* KO, *Slc35b*2 KO, and *B3galt1* KO samples had lower expression of *Dnmt3a* and *Dnmt3b*, mimicking the parental cell line cultured in 2i condition (Figure S3A). Interestingly, a few KOs led to notable repression of *Dnmt3a* (*Zbtb14* KO*, Mcm3ap* KO*, Spop* KO) and *Dnmt3b* (*Mcm3ap* KO*, Kmt2d* KO*, Bend3* KO), suggesting possible DNA methylation changes locally, at some *Dnmt3a/b*-targeted promoters.

Some germline genes are primarily regulated by DNA methylation; some by histone modifications and PRC complex; and some by both (Borgel et al., 2010; Hackett et al., 2012). A subset of these, including *Dazl* wild-type and mScarlet-inserted alleles, were also part of the 48 targets on the Fluidigm chip (Figure S3B). Unsurprisingly, the KO of known epigenetic regulators strongly induced all tested germline genes. Notably, *Kdm5c* KO cells highly upregulated almost all of them, confirming recent findings (Scandaglia et al., 2017), while *Bend3* KO cells induced the *Dazl* locus only (Figure S3B).

Finally, we measured the expression of several repeat elements families known to be regulated by various epigenetic pathways (Figure 3C). Intracisternal A-particles (IAP, IAPez) are induced in 2i condition (Figure S3A) and in the absence of DNA methylation (*Dnmt1* KO). They were also induced in *Kmt2d* KO and when any member of the TREX-2 complex was missing (*Mcm3ap* KO, *Sem1* KO, or *Eny2* KO). MLV (class I ERV) and MusD (class II ERV) elements were both upregulated in *Bend3* KO (Figure 3C). The other distinctive targets dysregulated by our KOs were MERVL and MMERVK10C (class III ERV): these were induced in the majority of KO populations, but not in 2i condition.

We then selected for further analysis 6 hits present in the different clusters that were not already known to regulate *Dazl*: BEND3, MCM3AP, KDM5C, KMT2D, SPOP, and ZBTB14 (Figure 3A-C, S3A). Among the 3 members of TREX-2 mRNA export complex, we chose to pursue MCM3AP since it is a scaffolding protein, important for the stability of the complex (Wickramasinghe et al., 2014) and it is also known to have additional nuclear functions (Eid et al., 2017; Evangelista et al., 2018).

### Generation and characterization of individual KO clones

Our previous results had been obtained on a heterogeneous population of cells expressing the KO sgRNAs. To verify these results and elaborate on them more precisely, we picked 3 individual KO clones for each of the 6 genes of interest. All clones had loss-of-function mutations in the targeted gene (Supplementary File 1) and survived Hygromycin selection. They reactivated mScarlet to various extents: while the control cells had ∼3% mScarlet-positive cells, *Mcm3ap* KO, *Spop* KO, *Zbtb14* KO, and *Bend3* KO clones had 20-30% mScarlet-positive cells, *Kdm5c* KO had ∼40%, and *Kmt2d* KO ∼90% (Figure 3D). The clones also expressed the untagged allele of *Dazl* at the RNA (Figure 3E) and protein level (Figure 3F). The degree of DAZL protein induction agreed with the percentage of mScarlet-positive cells, with *Kmt2d* and *Kdm5c* KOs causing the greatest DAZL induction (Figure 3F). The amount of DAZL expressed in serum condition by the KO clones was generally lower than the amount of DAZL expressed by 2i cells, showing that the induction is only suboptimal, or affects only a fraction of the cells, or both. The *Kmt2d* mutant clones deviated from this observation, and induced DAZL to near-2i levels (Figure 3F).

Western blot analysis of the clones for the expression of DNA methylation factors showed no change for UHRF1 (Figure S3C) or DNMT3A (differing from the transcript level for *Zbtb14* KO and *Mcm3ap* KO). Strikingly, the minor downregulation of *Dnmt3b* led to the loss of DNMT3B in *Kmt2d* KO clones (Figure S3C), which may explain the high level of expression of germline genes (DNMT3B being preferentially targeted at these loci (Velasco et al., 2010). RT-qPCR in the KO clones confirmed the transcriptional changes seen by Fluidigm on heterogeneous KO populations (Figure S3D).

### Rescue constructs revert the phenotypes of KO clones

We next verified that the phenotypes could be rescued by reintroduction of a WT cDNA; for this, we used a PiggyBac transposon-based system driving constitutive expression of V5-tagged proteins (Fukuda et al., 2018). We were not able to generate a rescue construct for *Kmt2d*, as the cDNA is very large (∼18kb), but we successfully expressed CRISPR-resistant alleles of mouse *Bend3*, *Kdm5c*, *Mcm3ap*, *Spop*, and *Zbtb14* in the corresponding KO lines. In all cases, expression of the functional gene decreased the number of mScarlet-positive cells to near-background levels (Figure 4A), rendered the cells sensitive to Hygromycin (Figure 4B), and silenced the expression of the *Dazl* RNA (Figure 4C) and protein (Figure 4D). In the case of KDM5C, we also tested rescue with a catalytically inactive mutant (H514A). The mutant was expressed to similar levels as the WT protein but was less effective than the WT protein at silencing mScarlet expression (Figure S4A) and Hygro^R^ expression (Figure S4B), However, endogenous allele is silenced by the catalytic dead KDM5C (Figure S4C). The effect of catalytic dead KDM5C at the two *Dazl* alleles, wild-type and reporter, was different indicating that the demethylase activity of KDM5C contributes to robust silencing but is not essential. These data prove that loss-of-function mutations in *Bend3*, *Kdm5c*, *Mcm3ap*, *Spop*, and *Zbtb14* have caused the reactivation of the epigenetic reporter used in our screen. The cells in which *Kmt2d* was targeted by sgRNAs contain loss-of-function mutations in the gene, and it is likely that these mutations underlie the epigenetic reactivation, but rescue experiments would be necessary to formally prove it.

**Figure 4:**
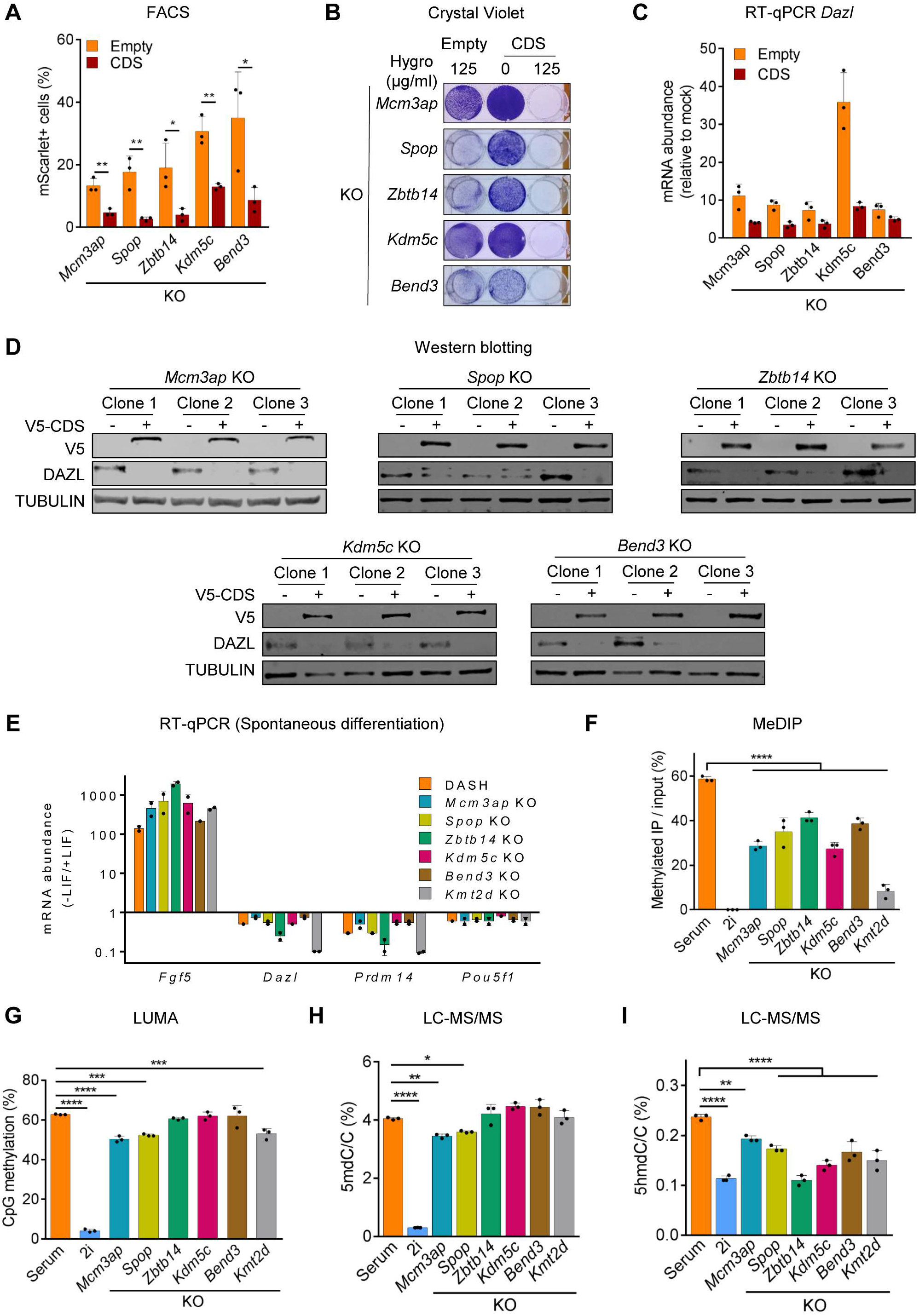
Rescue of Dazl expression in the 6 top hits. (A) mScarlet expression in the 6 indicated KOs, rescued with their respective V5-tagged protein-coding sequence (CDS), or with an empty vector. Each data point is an individual clone (n=3 clones per KO). (B) Hygromycin resistance after rescue. One representative clone is shown for each KO. (C) *Dazl* expression after rescue (n=3 rescued clones per KO). (D) Western blot analysis of DAZL and the indicated V5-tagged CDS. (E) RT-qPCR assays showing the expression changes of *Dazl*, differentiation (*Fgf5*) and pluripotency (*Prdm14, Pou5f1*) markers upon spontaneous differentiation after withdrawal of LIF. (F) MeDIP assay showing the relative levels of 5mC at the *Dazl* promoter for 6 candidates KO, compared to the parental DASH mES cells (n=3 clones per KO). (G) LUMA assay showing the changes in the global level of DNA methylation for 6 candidates KO, compared to the parental DASH mES cells (n=3 clones per KO). (H and I) LC-MS/MS assay showing the changes in the global level of 5mdC/C (H) and 5hmdC/C (I) for 6 candidates KO, compared to the parental DASH mES cells (n=3 clones per KO).

We then went on to characterize the cells with mutations in our 6 genes of interest. First, we verified more thoroughly that the cells were not unable to differentiate: each clone was left to undergo spontaneous differentiation by removal of LIF in the serum condition. In all cases, the cell colonies that were rounded in serum became irregularly shaped, which is a mark of differentiation; the *Tcf7l1* mutants, used as controls, failed to differentiate (Figure S4D). Pluripotency genes (*Prdm14*, *Pou5f1*) and *Dazl* were repressed, while differentiation marker *Fgf5* was induced (Figure 4E). These data are consistent with our previous RT-qPCR data and rule out the possibility that the KO cells are in a pseudo-2i state that is unresponsive to serum or differentiation cues.

The DNA methylation level at the *Dazl* promoter, assessed by MeDIP, was reduced in all KO clones but did not reach the very low level seen in 2i cells (Figure 4F). We also measured the global DNA methylation level, by LUMA and LC-MS/MS. The two approaches gave convergent results, with a small decrease of DNA methylation in *Mcm3ap* and *Spop* mutant cells, but no significant differences in the other mutants. Notably, we observed a decrease of 5hmdC level in all 6 KOs, suggestive of possible changes in the transcriptome considering the role of 5hmdC in transcription (Pastor et al., 2011) (Figure 4H). To better understand the phenomena leading to *Dazl* activation in the mutant cells, we turned to transcriptomic and DNA methylation analysis.

### Knock-outs display distinct transcriptomes

In a global DNA methylation analysis by WGBS, only *Spop* KO cells showed a significant genomic hypomethylation compared to parental cells (Figure S5A). However, a more precise analysis across specific genomic features revealed moderate changes in the KOs (Figure 5A-5B), indicative of localized regulation by DNA methylation in these cells. Next, we performed mRNA-seq on each of the mutant clones and compared their profile to cells grown in 2i or in serum. A Principal Component Analysis (PCA) yielded 3 important conclusions: 1) the mutant cells were very different from 2i cells, again ruling out a differentiation artifact; 2) the *Kdm5c* KO*, Mcm3ap* KO*, Spop* KO, and *Zbtb14* KO clones all clustered closely together, away from serum-grown control cells; 3) the *Bend3* and *Kmt2d* mutant cells each formed an isolated cluster, closer to the serum cells (Figure 5C). These data show that 4 mutants (*Kdm5c, Mcm3ap, Spop*, and *Zbtb14*) have similar characteristics and that they differ from *Bend3* and *Kmt2d* mutant cells.

**Figure 5:**
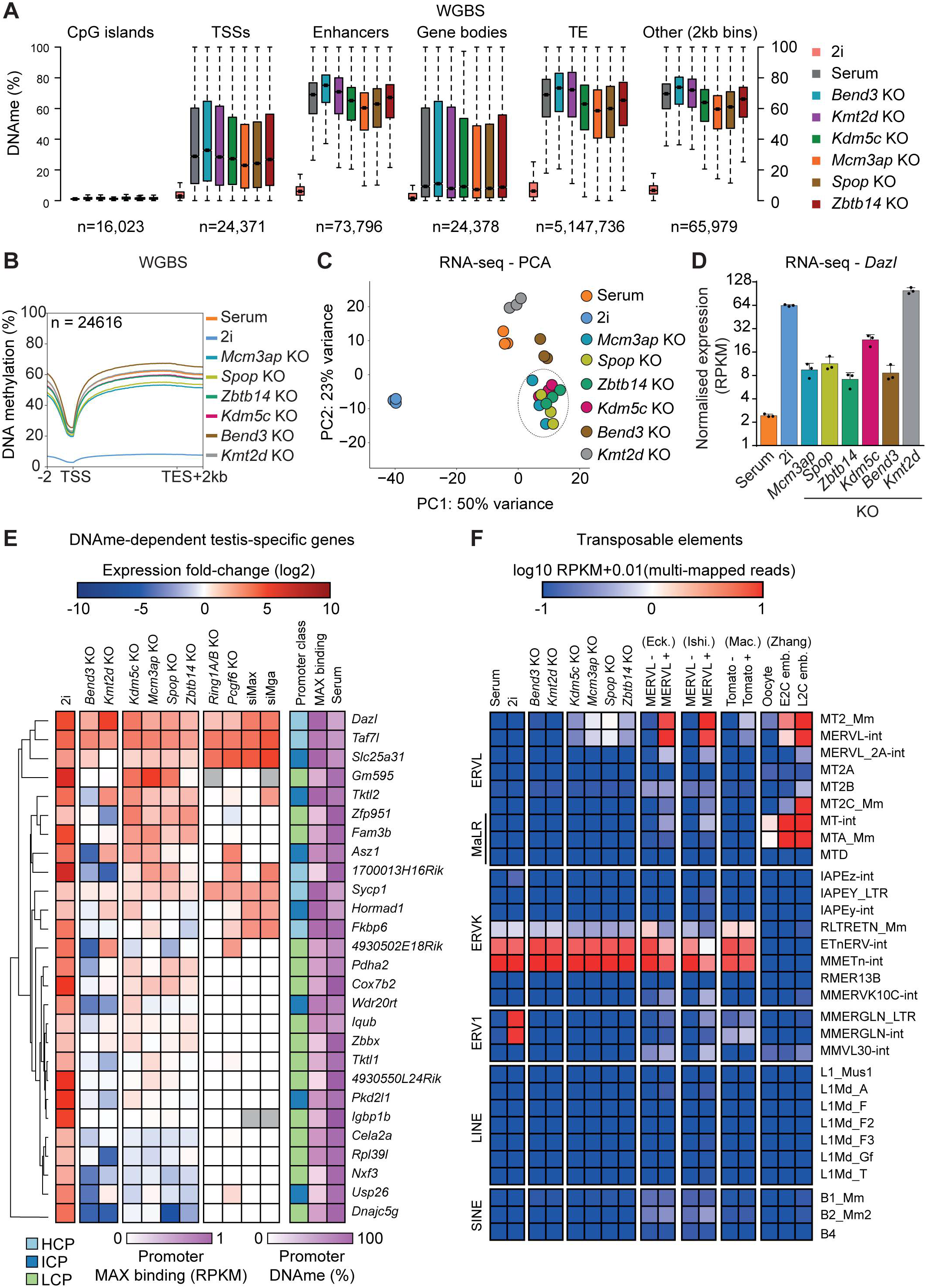
Knock-out of Spop, Mcm3ap, Kdm5c, and Zbtb14 induce similar transcriptomic perturbations. (A) Boxplots showing the distribution of CpG methylation levels in different genomic features in 6 candidates KO and parental DASH mES cells cultured in Serum or 2i. In the boxplots, the square indicates the median, the box limits indicate the upper and lower quartiles and the and whiskers denote the values lying within 1.5 times the interquartile range. (B) Metaplot of CpG methylation levels over RefSeq genes and 2 kb flanking sequences, calculated from the WGBS data for 6 candidates KO (n=3, clones per KO) and parental DASH mES cells cultured in Serum or 2i. (C) Principal Component Analysis (PCA) of the transcriptome (RNA-seq) of 6 candidates KO (n=3, clones per KO) and parental DASH mES cells cultured in Serum or 2i. (D) Normalized expression levels (RPKM) of *Dazl* in 6 candidates KO and parental DASH mES cells cultured in Serum or 2i. (E) Heatmap of the expression changes of the 27 DNA methylation-dependent testis-specific genes in 6 candidates KO and parental DASH mES cells cultured in 2i. The average log_2_(fold-change) relative to DASH mES cells cultured in Serum is shown for each sample. Published data for expression changes in the absence of ncPRC1.6 (*Ring1A/B* KO, *Pcgf6* KO, *Max* KD, *Mga* KD) was included for comparison. Promoters (TSS +/− 1kb) were categorized by CpG density (LCP, low CpG promoter; ICP, intermediate CpG promoter; HCP, high CpG promoter). Further, promoter DNA methylation levels in DASH mES cultured in Serum are shown. (F) Heatmap depicting the normalized expression (RPKM) of SINE, LINE, ERV1, ERVK, and ERVL retrotransposons in 6 candidates KO and parental DASH mES cells. Published data of MERVL-positive cells and 2C embryos were included for comparison. (A-F) For each candidate KO, n=3, clones per KO; and 3 biological replicates for parental DASH mES cells cultured in Serum or 2i.

To clarify these characteristics, we performed more specific transcriptome analyses and investigated Differentially Expressed (DE) genes. Using |FC| > 2 and *p*-adj < 1% thresholds, the KO cells showed from ∼1,000 (*Mcm3ap* KO) to ∼1,400 (*Kmt2d* KO) DE genes, roughly equally split between up- and down-regulated genes (Figure S5B), and as expected, *Dazl* was among the positively DE genes in all mutant lines (Figure 5D). Shared GO terms significantly enriched in up- or down-regulated DE genes included terms related to gene expression, cell differentiation, and metabolic processes (Figure S5C).

### Regulation of DNA methylation-dependent testis-specific genes

We next sought to determine whether these 6 factors were repressing loci similar to *Dazl,* a testis-specific gene. To this end, bioinformatic analysis was performed using public datasets to identify testis-specific genes regulated through epigenetic mechanisms, primarily by DNA methylation in mouse ESCs. We, therefore, analyzed RNA-seq datasets to identify mouse testis-specific genes (Figure S5D) and genes induced upon *Dnmt* triple KO, a complete absence of genomic DNA, along with differing contribution by inhibition of ncPRC1 complex using PRT4165 (Figure S5E). The transcriptome and methylome analysis of parental DASH mouse ESCs cultured in serum and 2i conditions identified genes whose expression was responsive to DNA methylation in mouse ESCs i.e. methylated and repressed in serum condition, but unmethylated and expressed upon switching to 2i condition (Figure S5F). The intersection of these datasets contained 27 DNA methylation-dependent testis-specific genes (hereafter referred to as “DMD-TSG”), including *Dazl* (Figure S5G). These genes have a similar expression profile as *Dazl,* thus considered if they have a similar or vastly different regulation from *Dazl.* However, many testis-specific genes that are primarily regulated by ncPRC1 in mouse ESCs were not considered for this analysis.

About one-third of the DMD-TSG are regulated by the ncPRC1 complex (*Ring1A/B, Pcgf6, Max, Mga*), reflecting the synergistic role it plays along with DNA methylation in gene repression, however, their contribution was minuscule compared to DNA methylation in mouse ESCs (Figure 5E). Of note, KDM5C repression showed the best overlap (6/7, 86%) with MAX-targeted genes. Moreover, the 4 knock-out lines (*Kdm5c* KO*, Mcm3ap* KO*, Spop* KO and *Zbtb14* KO) activated a similar set of genes: about half of the 27 DMD-TSG were upregulated upon KO, including many genes which are not regulated by ncPRC1 (Figure 5E). As expected, the gene activation was correlated with the reduced levels in average promoter DNA methylation, especially at *Dazl* promoter (Figure S5H).

Unlike the 4 clustered KOs, *Kmt2d* KO activated a smaller set of DMD-TSG whereas *Bend3* KO activated only 3 of them: *Dazl*, *Taf7l*, and *Slc25a31* (Figure 5E). Interestingly, *Kmt2d* KO and *Bend3* KO repressed several DMD-TSG which were activated in other KOs, highlighting a distinct gene regulation mechanism (Figure 5E). Notably, BEND3 activates several DMD-TSG (*Fkbp6, Hormad1*, *Nxf3*, *Usp26*, *Cela2a*, *4930502E18Rik*, *Rpl39l*), contrasting with its repressive function at *Dazl*. Intriguingly, KOs had heterogeneous effects on the expression of the genes also regulated by PRC1 (Figure 5E). For instance, *Fkbp6* and *Hormad1* are both induced by BEND3 and ZBTB14, but repressed by MCM3AP and KDM5C respectively. MCM3AP was also a common repressor of some PRC1-independent testis-specific genes, such as *1700013H16Rik* and *Asz1*. These differences among the 4 similar candidates suggest that they may have specific functions locally, depending on the chromatin context at these testis-specific gene promoters.

### Activation of repeat elements

We further analyzed our transcriptome data to determine whether the 6 genes under investigation regulated repeat elements (SINEs, LINEs, and ERVs). As documented previously, despite the global loss of DNA methylation in 2i, the majority of the repeat elements are repressed by alternative repressive pathways (Walter et al., 2016) (Figure 5F, S5I). We saw a notable induction of MERVL —but of no other repeats— in *Kdm5c* KO*, Mcm3ap* KO*, Spop* KO, and *Zbtb14* KO clones (Figure 5F). The expression of MERVL retroelements is normally restricted to the 2-cell stage of embryonic development; it is also expressed in 2C-like cells (2CLC) that appear spontaneously at low frequency when mouse ESCs are grown in serum (Macfarlan et al., 2012) (Figure 5F). While the transcriptional induction of MERVL repeats is clear, their level of DNA methylation assessed by WGBS is unaffected (Figure S5I), suggesting that a minority of repeats could be demethylated and active, or that the induction occurs in a minority of cells, or both.

### The KO of Kdm5c, Mcm3ap, Spop, or Zbtb14 induces a 2-Cell-like transcriptional signature

The 4 KOs have 201 common upregulated genes (Figure 6A) that are significantly expressed in 2C embryos compared to other early developmental stages (Figure 6B). Further, a GSEA and differential expression analysis of transcriptional profile against a list of 92 markers of the 2C-like state (Rodriguez-Terrones et al., 2018) showed a significant positive correlation for *Kdm5c, Mcm3ap, Spop* and *Zbtb14* KOs but not for *Bend3* and *Kmt2d* KOs (Figure 6C-D, S6A-B). Among the 2CLC signature genes, *Zscan4* cluster paralogs (*Zscan4a/b/c/d*/*f),* which are the predominant drivers of 2C state (Zhang et al., 2019), were upregulated in the 4 KOs but not in *Bend3* KO, *Kmt2d* KO or parental cells cultured in 2i condition (Figure 6D, S6B-C). Notably, upregulation of 201 common genes was not accompanied with any change in promoter DNA methylation (Figure S6D). Most of these genes are demethylated in 2i condition, yet are not expressed, highlighting alternative repressive mechanisms to keep them silent. Kock-out of *Kdm5c*, *Mcm3ap*, *Spop*, or *Zbtb14* possibly relieves this repression and/or requires expression of specific transcription factors.

**Figure 6:**
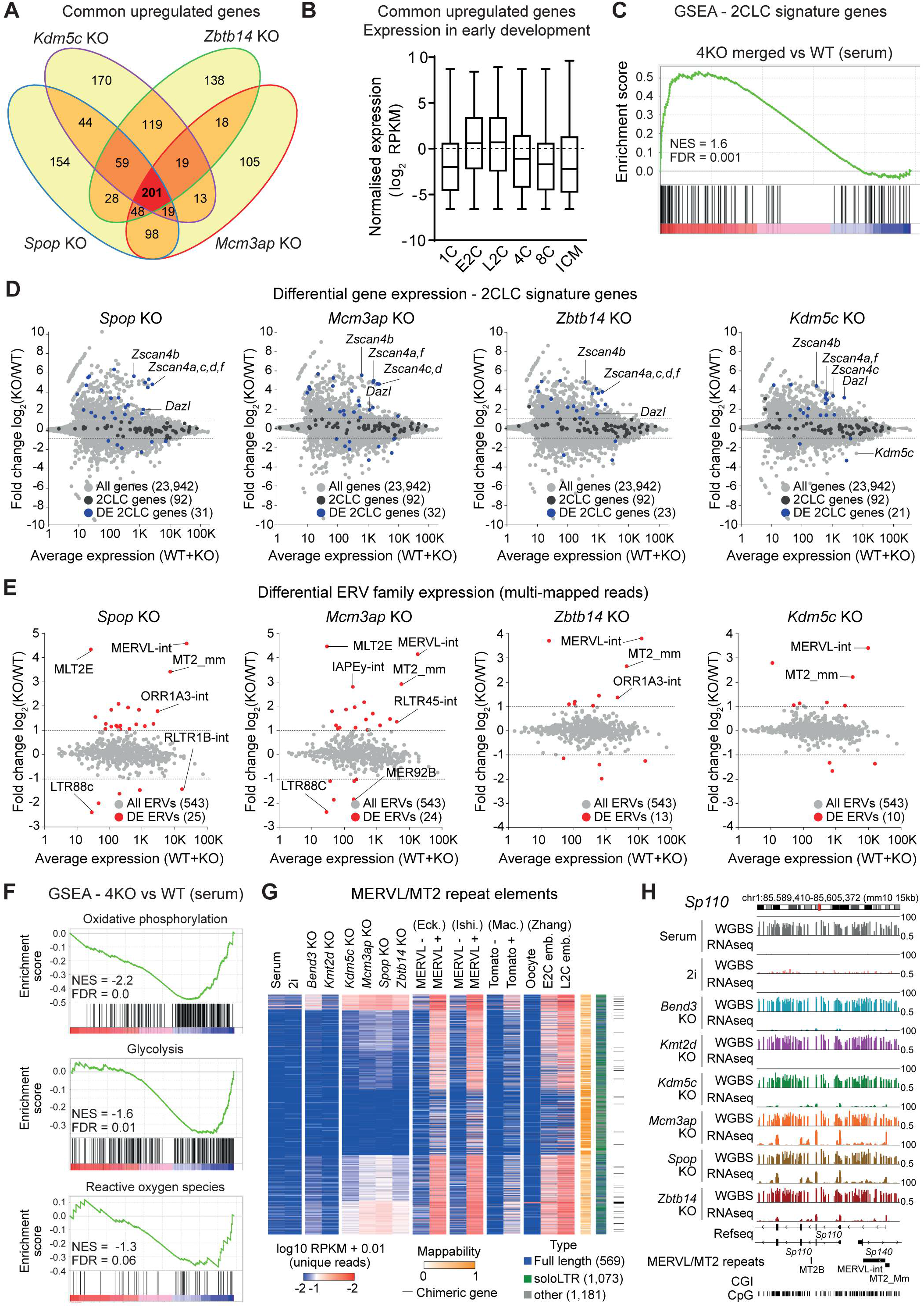
Knock-out of Spop, Mcm3ap, Kdm5c, and Zbtb14 induce 2CLC, totipotent-like state. (A) Venn diagram of upregulated genes (RNA-seq) after knock-out of *Spop*, *Mcm3ap*, *Kdm5c,* and *Zbtb14*. (B) Boxplots of normalized expression (RPKM) for 201 common upregulated genes in *Spop* KO, *Mcm3ap* KO, *Kdm5c* KO, and *Zbtb14* KO in early developmental stages. In the boxplots, the line indicates the median, the box limits indicate the upper and lower quartiles and the whiskers extend to 1.5 IQR from the quartiles. Transcriptome data was obtained from GSE66390 (Wu et al., 2016). (C) GSEA plot showing significant upregulation of 2CLC markers (Rodriguez-Terrones et al., 2018) in the 4 averaged transcriptome profiles (*Spop* KO, *Mcm3ap* KO, *Kdm5c* KO, and *Zbtb14* KO). (D) MA-plots showing differentially expressed 2CLC marker genes in *Spop* KO, *Mcm3ap* KO, *Kdm5c* KO, and *Zbtb14* KO cells. (E) MA-plots showing differentially expressed ERV retrotransposons in *Spop* KO, *Mcm3ap* KO, *Kdm5c* KO, and *Zbtb14* KO cells. (F) GSEA plots showing significant downregulation of selected metabolic pathways i.e., oxidative phosphorylation, glycolysis, and reactive oxygen species in the 4 averaged transcriptome profiles (*Spop* KO, *Mcm3ap* KO, *Kdm5c* KO, and *Zbtb14* KO). (G) Heatmap depicting the normalized expression (RPKM) of MERVL/MT2 single elements (full length, solo LTR, or others) in 6 candidates KO (n=3, clones per KO) and parental DASH mES cells. Mappability score and the presence of chimeric genes are indicated. Published data of MERVL-positive cells and 2C embryos were included for comparison and were obtained from GSE75751 (Eckersley-Maslin et al., 2016) E-MTAB-2684 (Ishiuchi et al., 2015) GSE33923 (Macfarlan et al., 2012) and GSE71434 (Zhang et al., 2016). (H) Genome browser tracks depicting WGBS and RNA-seq profiles at *Sp110* locus in 6 candidates KO and parental DASH mES cells cultured in Serum or 2i (RNA-seq tracks and WGBS of KO cells correspond to the average of 3 replicates). Chimeric transcript originating from MERVL/MT2 repeats is indicated.

### Knock-outs induce 2CLC metabolic state and MERVL-LTR chimeric transcripts

Another feature of 2CLC is a metabolic switch with marked changes in oxidative metabolism: reduction of oxygen consumption, glycolytic activity and ROS accumulation (Rodriguez-Terrones et al., 2020). GSEA analysis employing curated Hallmark pathways showed that ‘Oxidative phosphorylation’, ‘Glycolysis’ and ‘Reactive Oxygen Species’ were among the top significantly depleted pathways for the combined 4KO profile, but not for *Bend3* KO, *Kmt2d* KO and parental cells cultured in 2i condition (Figure 6F, S6E), emphasizing a specific induction of 2CLC state. Previously, we observed the expression of MERVL repeated elements, a marker of the 2CLC state. We performed a differential ERV family expression analysis, detected an upregulation of MERVL and LTR in the 4 KO, a subtle induction in *Bend3* KO, no derepression in *Kmt2d* KO and parental cells cultured in 2i (Figure 6E, S6F-G). Further, DNA methylation-dependent IAPEy elements which are expressed in 2i, were uniquely induced in *Mcm3ap* KO (Figure 6E, S6H).

During zygotic genome activation (ZGA), many of the MERVL-LTR drive the expression of ZGA genes (Eckersley-Maslin et al., 2019). In this network, *Dux* is a master regulator of 2C state creating a feedback loop to drive the expression of ZGA genes until its deactivation by LINE-1 elements, permitting the transition to 4C cell stage (Percharde et al., 2018). Indeed, we observed the expression of *Dux* in the 4 KOs (Figure S6I). MERVL elements have thousands of copies across the genome, ranging from full length LTR spanning >6000bp with both LTR promoters and internal sequences, MT2_Mm LTR promoter elements under 500 bp in length referred as “Solo-LTRs” and other truncated versions (Macfarlan et al., 2012; Morgani et al., 2013). Detailed analysis on unique MERVL/MT2 reads showed that only a subset of 2C activated copies were upregulated and were very similar in the 4 KO (Figure 6G). This subset did not depend on the mappability of the reads (reflecting their divergence from the consensus sequence) but was clearly induced in previously characterized 2CLC (Eckersley-Maslin et al., 2016; Ishiuchi et al., 2015; Macfarlan et al., 2012). Many of these upregulated MERVL LTR copies led to the expression of chimeric transcripts, such as *Sp110* (Figure 6H), known to positively regulate the 2CLC state (Eckersley-Maslin et al., 2019).

### Rescue of 2CLC phenotype

We validated RNA-seq results by RT--qPCR for 2C markers (Figure 7A). Compared to parental DASH mouse ESCs, 4 KOs showed a strong induction of *Dux,* MERVL and *Zscan4d*. We also validated the findings at the protein level and observed induction of ZSCAN4 and MERVL-Gag in the KO clones for *Kdm5c*, *Mcm3ap, Spop*, and *Zbtb14*, but not for *Bend3* and *Kmt2d* KOs (Figure 7B, S7B). Importantly, the expression of *Zscan4* at the RNA and protein levels was abolished when the mutations were rescued by the corresponding WT cDNAs (Figure 7C-D, S7C). Intriguingly, the induction of 2C markers was also completely rescued by KDM5C catalytically inactive mutant (H514A) (Figure S7D-E), meaning that the histone demethylase function is dispensable in this context (Figure S4A-C). In summary, our system based on *Dazl* is sensitive to epigenetic perturbations and useful to report on repressive pathways in mouse ESCs. We used this system for a genome-wide CRISPR knock-out screen, which yielded 6 novel epigenetic regulators: ZBTB14, KDM5C, SPOP, MCM3AP, BEND3, and KMT2D. All the identified 6 candidates regulate the expression of germline genes, however, they act on a different set of germline genes. Further, kock-out of ZBTB14, KDM5C, SPOP and MCM3AP led to reactivation of the 2CLC state.

**Figure 7:**
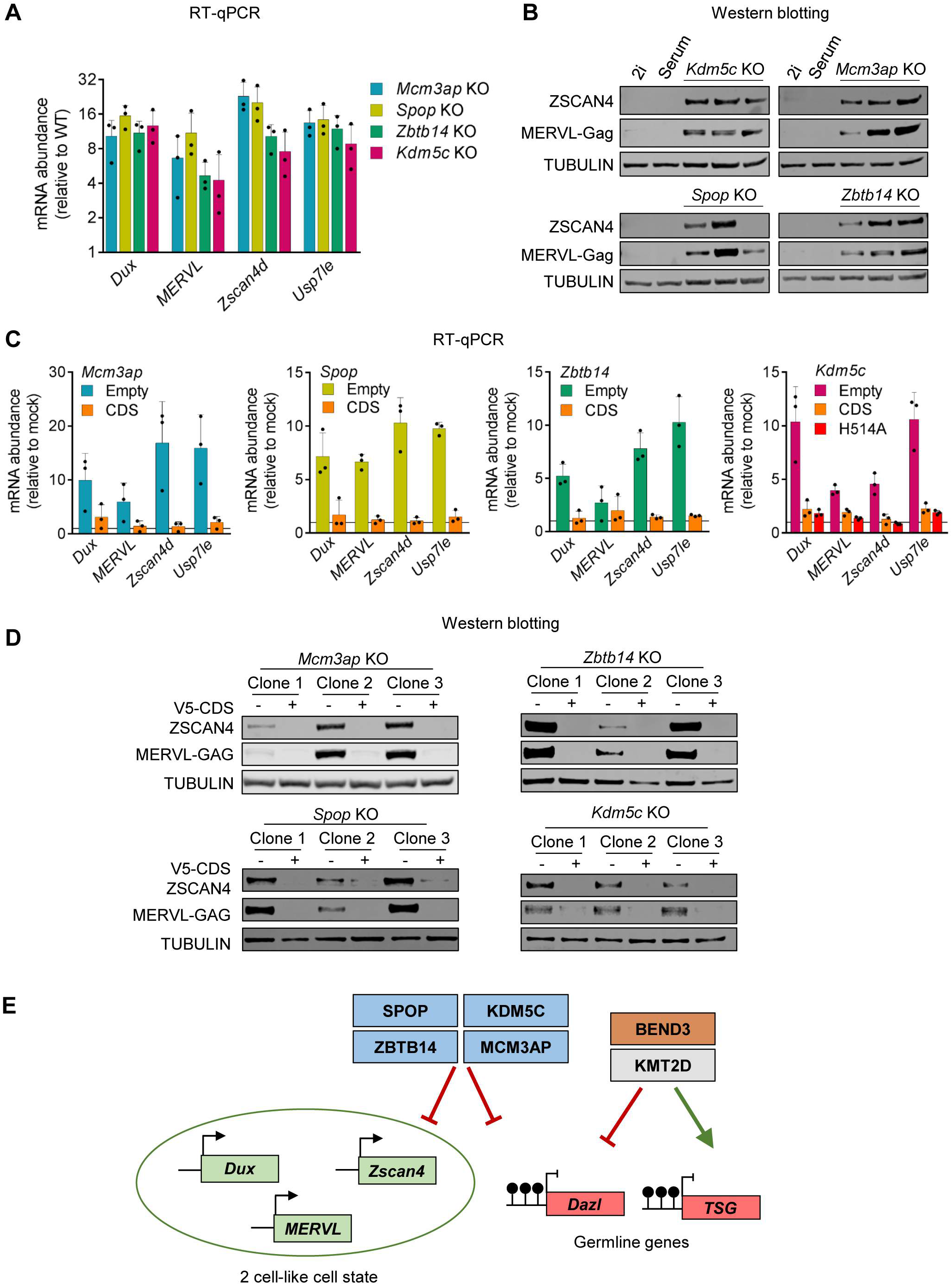
Rescue of 2CLC phenotype in 4 novel candidates KO. (A) RT-qPCR analysis of the expression of 2C-specific genes in *Spop* KO, *Mcm3ap* KO, *Kdm5c* KO, and *Zbtb14* KO, compared to the parental DASH mES cells (n=3 clones per KO). (B) Western blot analysis of ZSCAN4 and MERVL-Gag for *Spop* KO, *Mcm3ap* KO, *Kdm5c* KO, *Zbtb14* KO, and parental DASH mES cells cultured in Serum or 2i. (C) RT-qPCR analysis of the expression of 2C-specific genes for *Spop* KO, *Mcm3ap* KO, *Kdm5c* KO, and *Zbtb14* KO, transfected with either an empty plasmid or their respective CDS, compared to the parental DASH mES cells (n=3, clones per KO). (D) Western blot analysis of V5, ZSCAN4, and MERVL-Gag in 4 candidates KO, transfected with an empty plasmid or their respective V5-tagged CDS. (E) Model for role of novel epigenetic factors ZBTB14, KDM5C, SPOP, MCM3AP, BEND3, and KMT2D in the regulation of germline genes, and 2 cell-like cell state.

## Discussion

### DASH, an ideal reporter to study epigenetic repressive pathways

In our search for new actors in the epigenetic repression machinery, we performed a genome-wide CRISPR knock-out screen with a positive selection for cells re-expressing the DASH reporter, comprising of two selectable markers (for fluorescence, mScarlet, and antibiotic resistance, Hygro^R^), inserted in-frame of *Dazl* through a targeted CRISPR knock-in. DAZL is uniquely expressed in the germline lineage and during the wave of mouse ZGA, thus establishing *Dazl* as a marker for germ cells and 2C developmental stage. *Dazl* repression in other developmental stages is epigenetically mediated including promoter DNA methylation and PRC1 complex.

Our reporter DASH differs from previously published DNA methylation-sensitive reporter systems based on *Dazl* promoter as a DNA methylation sensor (Stelzer et al., 2015; Gretarsson & Hackett, 2020) as it is anchored in an endogenous genomic context allowing the study of multiple epigenetic pathways regulating its endogenous expression. Thus, the expression of these two markers in DASH depends on the regulation of *Dazl* expression, activated upon relieving the repression by DNA methylation, PRC complex, repressive histone post-translational modifications (PTMs), potentially assisted by the presence of transcription factors and activating histone PTMs.

The screen successfully recovered known regulators of *Dazl* such as DNMT1, UHRF1, and MGA but also several novel epigenetic regulators. One key limitation of our experimental procedure is that KOs in genes that are essential or near-essential in ES cells are strongly counter-selected in the course of the antibiotic selection. This likely explains why MAX, which is a well-validated repressor of *Dazl* (Endoh et al., 2017; Stielow et al., 2018), has a poor ranking in our list of hits (#1064). Possible avenues to capture some of the missing factors in the future would be to perform fluorescence-based screens and to examine cells at earlier time points.

Notwithstanding these limitations, our screening procedure was robust, as most novel hits (15/18, 83%) could be re-validated by generating new KOs; also, in all the cases we tested, we could rescue the KO phenotypes by expressing the corresponding sgRNA-resistant WT cDNA. We validated and characterized novel epigenetic regulators of *Dazl:* SPOP, MCM3AP, ZBTB14, KDM5C, BEND3, and KMT2D. This means that our procedure could be used immediately for further screens; those could be genetic using CRISPRi targeting lncRNAs, for instance, but chemical screens could also be performed.

### Regulation of DNAme-dependent testis-specific genes in mES cells

*Dazl* plays an important role in germ cells and potentially shares a similar regulation with other germline genes in mouse ESCs. By bioinformatic analysis, we identified 27 testis-specific genes that are upregulated in mouse ESCs in the absence of a repressive chromatin landscape i.e., in *Dnmt*-TKO and 2i conditions. In mouse ESCs, *Taf7l* was the only gene induced by all the KOs, however, its expression has been shown to depend on *Dazl* during germ cell development (Gura et al., 2020), a similar feature could be driving its expression in mouse ESCs. Besides this, all 6 gene KOs activated a different set of DNAme-dependent testis-specific genes in mES cells, some shared, some unique.

In line with the PCA of transcriptomic data, knock-out of the 4 clustered candidates (*Spop, Zbtb14, Mcm3ap,* and *Kdm5c*) led to the induction of a similar subset of testis-specific genes. Most of the testis-specific genes synergistically silenced by DNA methylation and MAX/MGA were upregulated in any of the 4 KOs. However, some genes not repressed by ncPRC1.6 (eg *Asz1*, *Fam3b, Pdha2)* were also induced upon KO, possibly indicating that they can regulate targeting of ncPRC1 and DNA methylation machinery on these loci and/or act at these loci directly. Notably, KDM5C has previously been reported to be part of a complex comprising several ncPRC1.6 subunits (MAX, MGA, DP1, E2F6, HDAC1/2) (Tahiliani et al., 2007). KDM5C has also been shown to be essential to silence germline genes during neuronal maturation (Scandaglia et al., 2017).

*Kmt2d* KO and *Bend3* KO, on the other hand, activated very few testis-specific genes and repressed several others, indicating that they may play a very direct role at these loci. KMT2D is an H3K4 methyltransferase, targeting enhancers (Wang et al., 2016). KMT2D positively regulates the expression of *Dnmt3b* (Cao et al., 2018; Dhar et al., 2018) and we observed reduced levels of DNMT3B upon *Kmt2d* KO. Further, DNMT3B is the primary DNAme regulator for *Dazl* (Velasco et al., 2010) and indeed DNA methylation level was dramatically decreased at *Dazl* promoter in *Kmt2d* KO cells, however, globally levels were not perturbed. BEND3 is known to function as a transcriptional repressor on heterochromatin (Sathyan et al., 2011), where it recruits PRC2 and NuRD complexes at unmethylated major satellites (Saksouk et al., 2017). Here, we demonstrate a role for BEND3 on the protein-coding genes, as a repressor of *Dazl* but also as an activator of multiple DNA-methylation-dependent testis-specific genes. Interestingly, local DNA methylation level decreased at *Dazl* promoter in *Bend3* KO cells, however, for many other induced testis-specific genes, DNA methylation level was unchanged. This exemplifies potential roles for *Bend3* as both a repressor and an activator at protein-coding genes.

This screen based on *Dazl*, highlights the diverse regulation of testis-specific genes in mouse ESCs and utility of germline genes to conduct similar screens to identify additional epigenetic regulators. A deeper understanding of the regulation of repressive pathways regulating testis-specific genes will be critical to understanding the activation of cancer-testis antigens seen in many cancers and their utility as a marker for diagnostic and prognostic purposes and also for potential therapeutics (Gjerstorff et al., 2015; Naciri et al., 2019).

### Regulation of 2CLC state

In addition to being a testis-specific gene, *Dazl* is a marker of the 2CLC state. A common feature, seen among the 4 KOs (*Spop, Zbtb14, Mcm3ap,* and *Kdm5c*) was the emergence of a particular cellular state i.e., totipotent-like 2 cell-like cells, sharing properties and markers with the 2-cell embryo, which mirrors in culture the transition from maternal to zygotic transcriptional programs occurring during mouse early development (Eckersley-Maslin et al., 2018). Over the last decade, several epigenetic factors activating or inhibiting the 2CLC state have been identified (Iturbide & Torres-Padilla, 2020), including non-coding RNAs (Choi et al., 2017; Percharde et al., 2018; Yang et al., 2020).

Several known negative regulators of the 2CLC state were present among the top hits of the DASH screen. For instance, the SUMO E3-ligase PIAS4 ranked #41 (Yan et al., 2019); the chromatin-remodeler p400 ranked #52 (Rodriguez-Terrones et al., 2018); the histone demethylase LSD1 ranked #111 (Macfarlan et al., 2012); the ncPRC1 subunits RING1B and RYBP ranked #100 and #245 respectively (Rodriguez-Terrones et al., 2018). However, many of the previously known 2CLC epigenetic regulators such as histone chaperone CAF-1 (Ishiuchi et al., 2015), transcription factor TRIM28 (De Iaco et al., 2017), the H3K9 methyltransferase SETDB1 (Wu et al., 2020), and the m^6^A binding protein YTHDC1 (Liu et al., 2021) were not ranked high in our screen. This could be due to the lethality associated with these KOs, especially considering the timeline of the DASH screen (3 weeks) in comparison to siRNA screens (Rodriguez-Terrones et al., 2018) with short timelines, where these factors were identified. In addition, some of the previously characterized regulators, which induced the expression of MERVL-LTR-based reporters (Rodriguez-Terrones et al., 2018), perhaps do not induce *Dazl* expression.

Compared to ES cells cultured in serum, 2CLCs have a distinct transcriptome profile (Rodriguez-Terrones et al., 2018), more accessible chromatin (Genet & Torres-Padilla, 2020), reduced DNA methylation (Eckersley-Maslin et al., 2016; Dan et al., 2017), fewer repressive marks (Eckersley-Maslin et al., 2018) and exhibit a change in oxidative metabolism (Rodriguez-Terrones et al., 2019). We were able to capture upregulated 2CLC signature at the transcriptional level and downregulation of oxidative metabolic pathways upon the KO of novel epigenetic repressors of the 2CLC state. Since only a small percentage of these cells exist in the 2CLC state, changes in the DNA methylation are harder to ascertain by the WGBS data obtained from the total cell population.

During evolution, mammalian genomes have accumulated a substantial amount of transposable elements in their genome. The silencing of these repeat elements is critical for the maintenance of genome integrity (Jacobs et al., 2014), and their misexpression and/or genome hopping is often related to cancer (Jang et al., 2019). At the same time, many of the repeat elements have coadapted and are essential for development (Rowe & Trono, 2011). During early development, at the 2C embryo stage, expression of the MERVL retrotransposons and associated LTRs takes place in this specific window. A master regulator of the 2CLC transcriptome is the transcription factor DUX, which binds and activates MERVL repeats and many promoters in this program and is instrumental for the zygotic genome activation (De Iaco et al., 2017). The exit from 2C to 4C stage is determined by another repeat element, LINE1, acting as a scaffold for recruiting *Dux* repressors (Percharde et al., 2018).

Accordingly, all 4 KOs strongly induced MERVL repeats, associated LTRs, and *Dux* which drive the 2-cell transcriptional program (Hendrickson et al., 2017; Whiddon et al., 2017). Notably, the expression of LINE1 repeats was unchanged in the total cell population. However, it remains to be investigated if LINE1 plays a similar role in the repression of *Dux* and MERVL in 2CLC. This is particularly challenging as 2CLC do not have a documented self-renewal capacity (Eckersley-Maslin et al., 2016)

Additionally, *Mcm3ap* KO activated another family of retrotransposons: IAPey elements. These are usually repressed in mESCs by multiple pathways: by DNA methylation and H3K9me3 (He et al., 2019), by H3K9me3/H3K27me3 in the absence of DNA methylation (Walter et al., 2016), by endosiRNA (Berrens et al., 2017) and by m^6^A RNA methylation as shown recently (Chelmicki et al., 2021). Considering the specific effect on IAPey elements in *Mcm3ap* KO, we speculate that MCM3AP could either act directly by binding at these loci or through regulation of another factor (through its TREX-2 RNA processing) acting uniquely at these loci, as a part of several of these repressive pathways.

### Possible mechanisms for 2CLC reactivation after KO

The role of *Dux* in mouse embryogenesis is debatable, upon its KO, there are developmental defects, nevertheless, pups are viable (Chen & Zhang, 2019; De Iaco et al., 2020). However, in the context of 2CLC, the majority of the known epigenetic regulators act via *Dux* and indeed was activated upon the knock-out of identified 4 novel 2CLC repressors. *Dux* is expressed in a specific developmental window, ectopic expression of *Dux* and its downstream targets in muscle tissue leads to facioscapulohumeral dystrophy (FSHD) (Tassin et al., 2012; Young et al., 2013). Mutations in CAF-1 and DNMT3B (Campbell et al., 2018; van den Boogaard et al., 2016), two known epigenetic factors repressing *Dux,* are associated with this disease. A similar loss of repression mechanisms could explain *Dux* induction in *Kdm5c* KO cells via the interaction between KDM5C and the ncPRC1.6 complex (Rodriguez-Terrones et al., 2018).

Conversely, some KOs may activate pathways promoting *Dux* induction. For instance, ATR-dependent replication stress triggers *Dux* expression (Atashpaz et al., 2020) and the zinc-finger protein ZBTB14 has been recently shown to stabilize the RPA-ATR-ATRIP complex at stalled replication forks (Kim et al., 2019). Loss of ZBTB14 can potentially trigger replication stress, leading to *Dux* expression. SPOP is an E3-ubiquitin ligase promoting the proteasome-mediated degradation of a wide range of substrates (Clark & Burleson, 2020), including DPPA2 (Rolland et al., 2014). DPPA2 binds directly at the *Dux* locus and activates it (Eckersley-Maslin et al., 2019). Interestingly, DPPA2 also binds to the *Dazl* promoter (Gretarsson & Hackett, 2020). Thus, the 2CLC state may arise from the loss of SPOP-dependent degradation of DPPA2, as observed in testicular germ cell tumors (Wang et al., 2021).

### Potential avenues for 2CLC for gene and cell therapeutics

Contrary to the pluripotent stem cells that can be potentially transformed into any adult somatic or germ cells, 2CLCs are believed to be very close to 2C cells, that are totipotent, i.e., able to form an entire organism by itself without carrier cells (Riveiro & Brickman, 2020). However, designing robust assays to assess this characteristic remains challenging, as current assays such as chimera formation and nuclear transfer assays do not directly inform on totipotency potential (Genet & Torres-Padilla, 2020). Another important experimental issue is the seeming lack of self-renewal capacity for 2CLC, thus limiting their use for cell and gene therapeutics. Gaining insights into the 2CLC regulatory network may contribute to overcome this technical problem and allow a stable culture of 2CLC and/or controlled cell state transition. Then, similarly to ESC reprogrammed to iPSC (Takahashi & Yamanaka, 2006) or EPSC (Yang et al., 2017), might be able to develop 2CLC use for cell therapeutics purposes.

## Supplemental Figure legends

**Figure S1:**
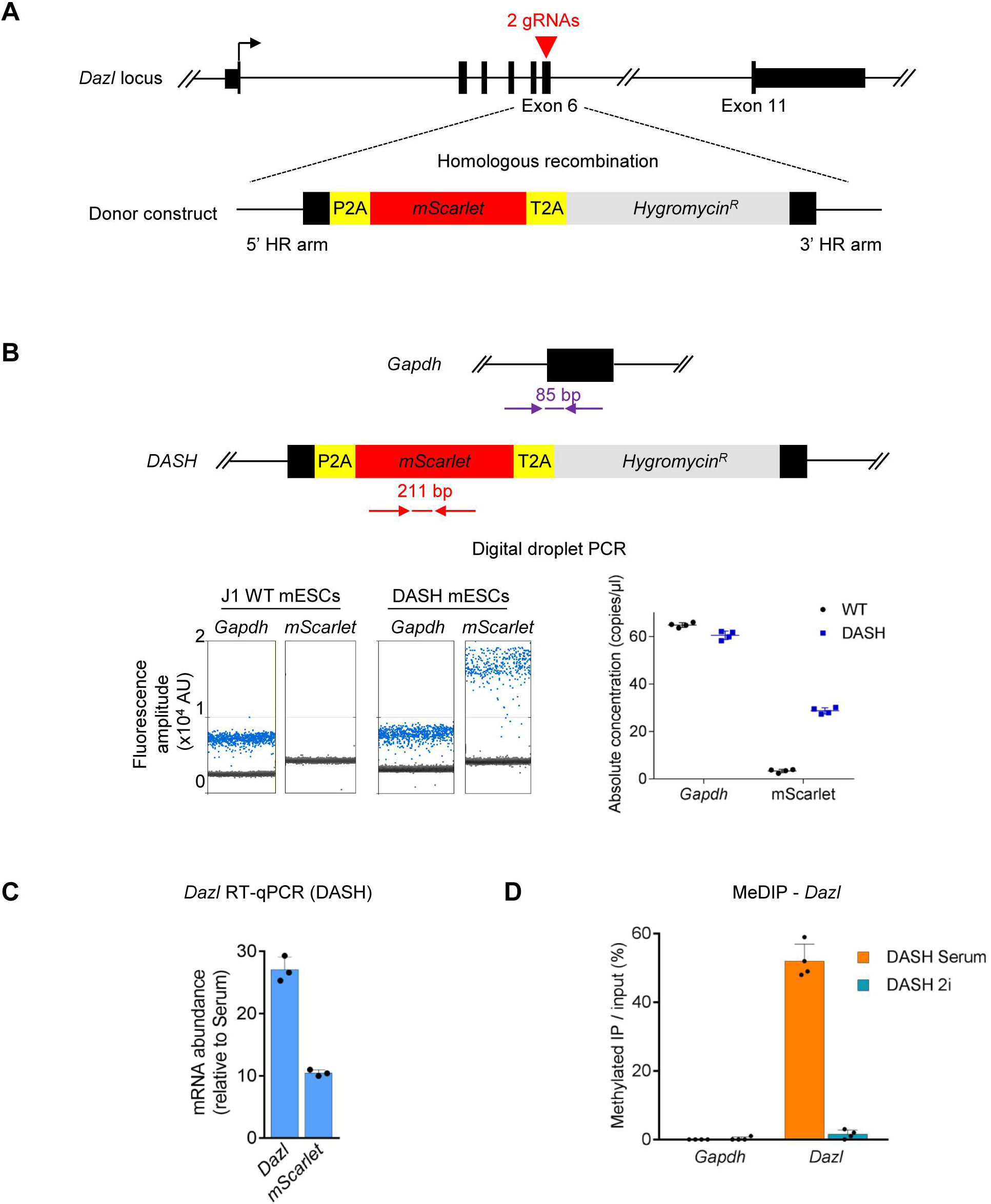
Generation and validation of Dazl-mScarlet-Hygromycin (DASH) reporter cell line. (A) *Dazl* exon 6 was targeted by 2 independent sgRNAs (red arrowheads) to insert the reporter cassette by homologous recombination. The donor construct contains *Dazl* homology arms flanking genes for the red-fluorescent protein mScarlet, and the Hygromycin resistance enzyme (HygroR) separated by 2A self-cleaving peptides (P2A, T2A). (B) DASH mES cells have a single insertion at one of the *Dazl* alleles, as determined by ddPCR. Top panel: schematic of primer pairs used for ddPCR. *Gapdh* served as controls present at 2 copies/cell. Bottom left panel: Blue droplets are positive for the corresponding amplification; black droplets are negative. About 18,000 droplets were analyzed for each amplification. Bottom right panel: quantitative analysis confirming single insertion of the donor construct. (C) RT-qPCR showing the up-regulation of *Dazl* and *mScarlet* in DASH mES cells cultured in 2i (relative to Serum condition). (D) MeDIP assay showing the relative levels of 5mC at *Gapdh* and *Dazl* promoters in DASH mES cells cultured in Serum and 2i conditions.

**Figure S2:**
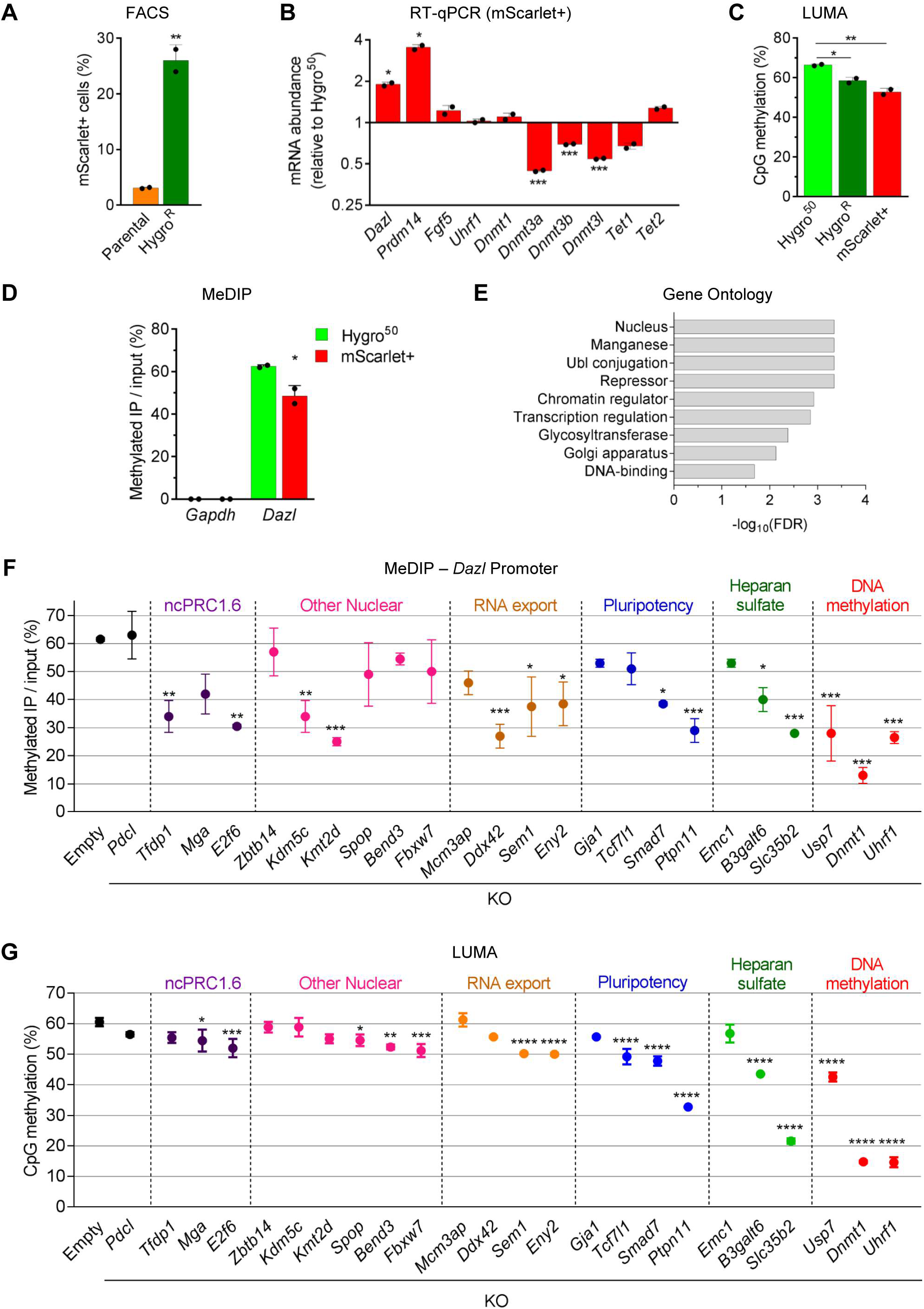
Knock-out of identified candidate genes in DASH mES cells validates CRISPR KO screen results. (A) FACS analysis of mScarlet expression after Hygromycin selection (the 2 replicates of the screen are shown). (B) RT-qPCR: comparison of mScarlet-expressing cells to Hygro^50^ cells. (C) Decrease of global DNA methylation in mScarlet+ cells in comparison to Hygro^50^ cells from the screen, as measured by LUMA. (D) MeDIP assay showing the relative levels of 5mC at *Gapdh* and *Dazl* promoters in Hygro^R^ and mScarlet+ screen samples. (E) Gene ontology (GO) terms (Uniprot keyword) significantly enriched among the top 40 hits. (F) MeDIP assay showing the relative levels of 5mC at *Dazl* promoter after KO of the indicated candidates and Hygromycin selection. (G) LUMA assay showing the changes in the global level of DNA methylation after KO of the indicated candidates and Hygromycin selection. For both MeDIP and LUMA, cells transduced with an empty vector (“Empty”) served as a negative control.

**Figure S3:**
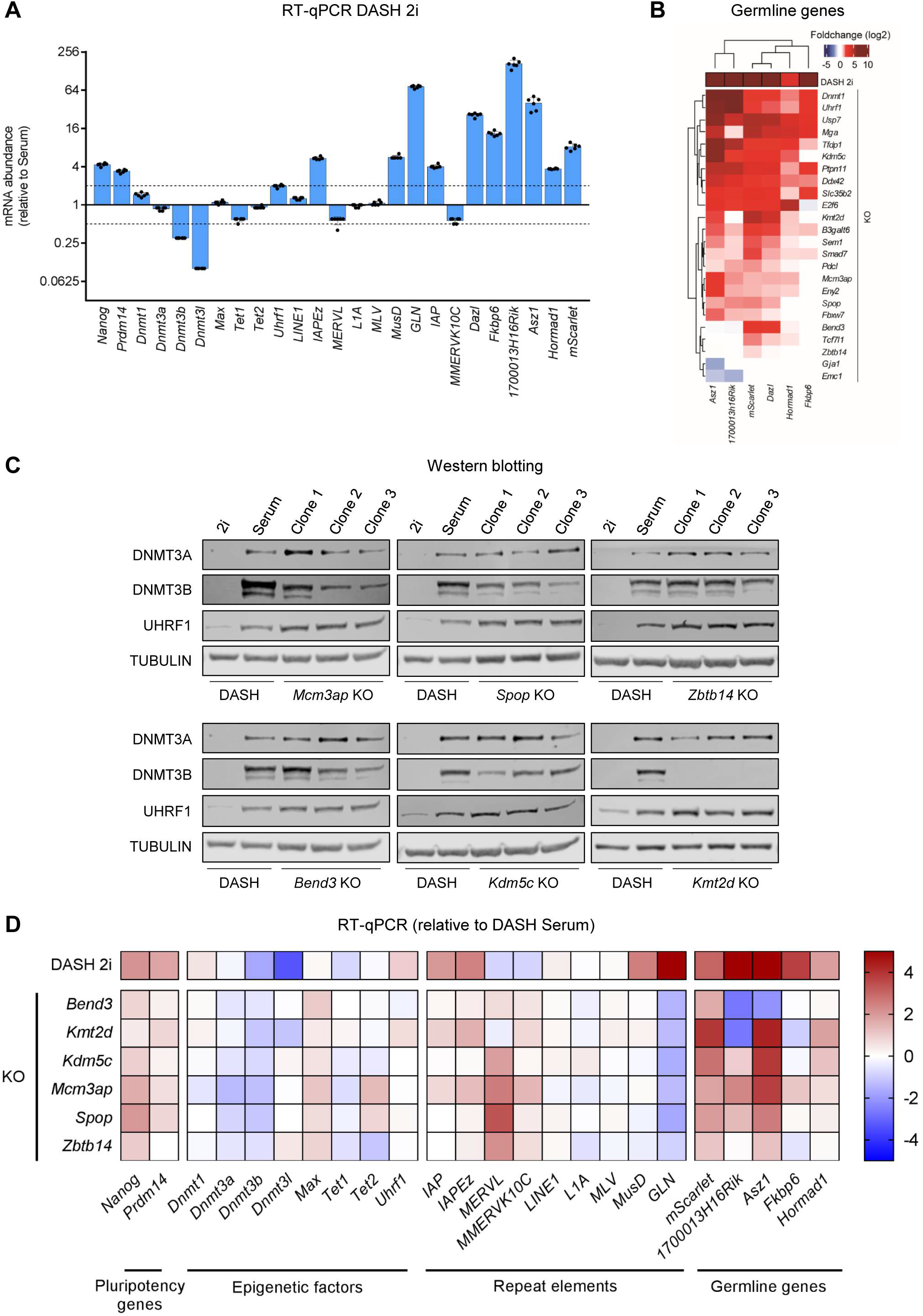
Novel epigenetic regulators of Dazl do not incur global changes in cell-state or epigenetic state. (A) RT-qPCR assays showing the expression changes of genes tested in the Fluidigm assay, in DASH mES cells cultured in 2i condition (relative to Serum). (B) Hierarchically clustered heatmaps from Fluidigm assay of Hygromycin-resistant individual candidate KO lines showing effects on the expression of germline genes. (C) Western blot analysis of DNMT3A, DNMT3B, and UHRF1 in parental DASH mES cells (cultured in Serum or 2i), and Hygromycin-resistant candidate KO clones. (D) Heatmap for RT-qPCR assay showing the expression changes of genes tested in the Fluidigm assay, in Hygromycin-resistant candidate KO clones. For each sample, the average log_2_(fold-change) value of 3 clones is depicted, relative to DASH mES cells cultured in Serum.

**Figure S4:**
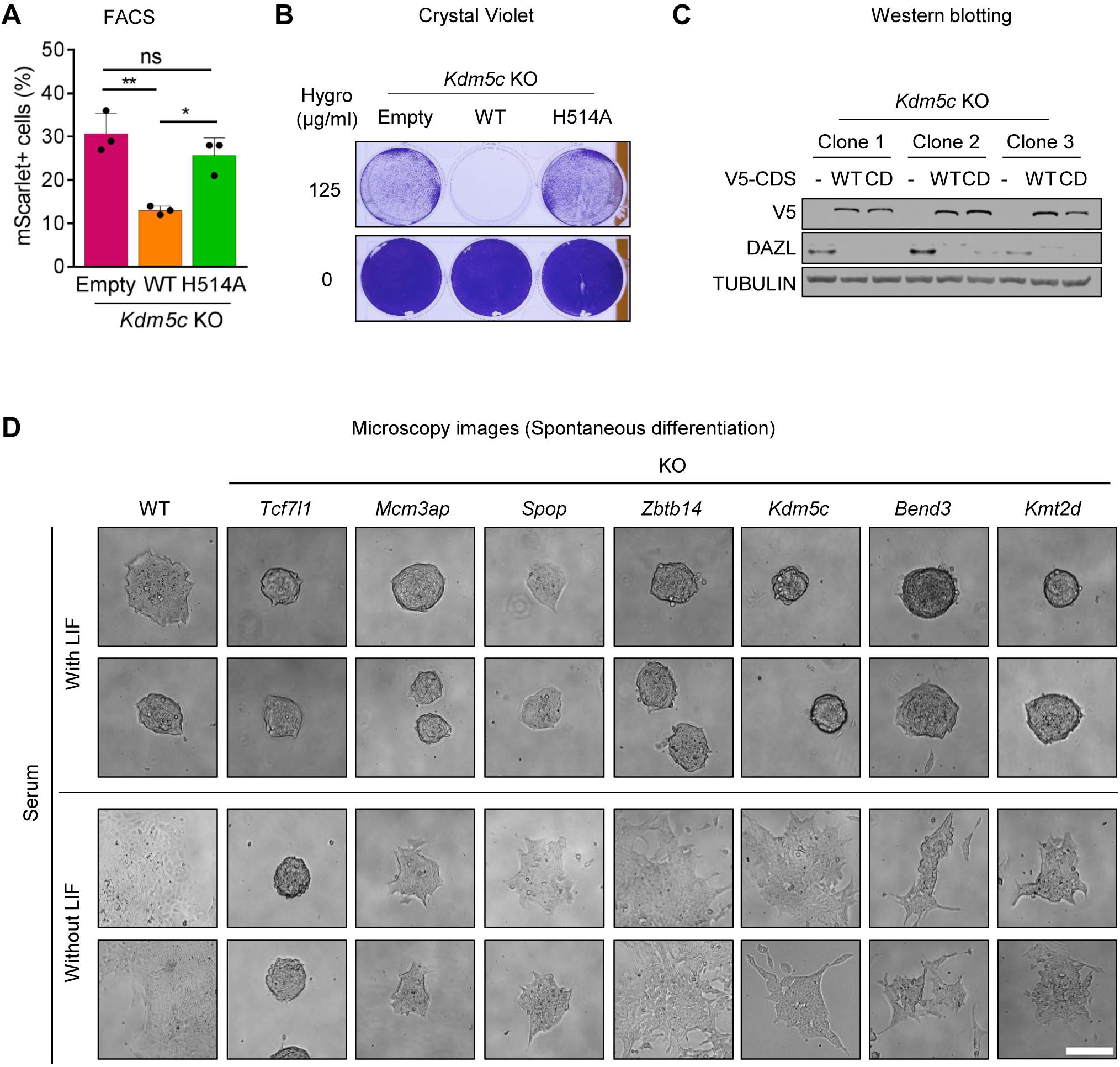
Novel epigenetic regulators of Dazl are not locked in a naïve pluripotent state. (A-C) *Kdm5c* KO clones were transfected with either an empty plasmid, or a V5-tagged wild-type *Kdm5c* CDS, or a V5-tagged catalytically-dead *Kdm5c* CDS (H514A) and were analyzed by Flow cytometry for mScarlet expressing cells (A), for cell survival by crystal violet staining upon Hygromycin selection (B), and by Western blot for DAZL and V5 protein expression. (D) Representative microscopy DIC images of KO mES colonies cultured in Serum, with LIF (control) or without LIF (spontaneous differentiation). *Tcf7l1* KO served a negative control as mES cells can’t differentiate without TCF7L1. Scale bar: 200 µm.

**Figure S5:**
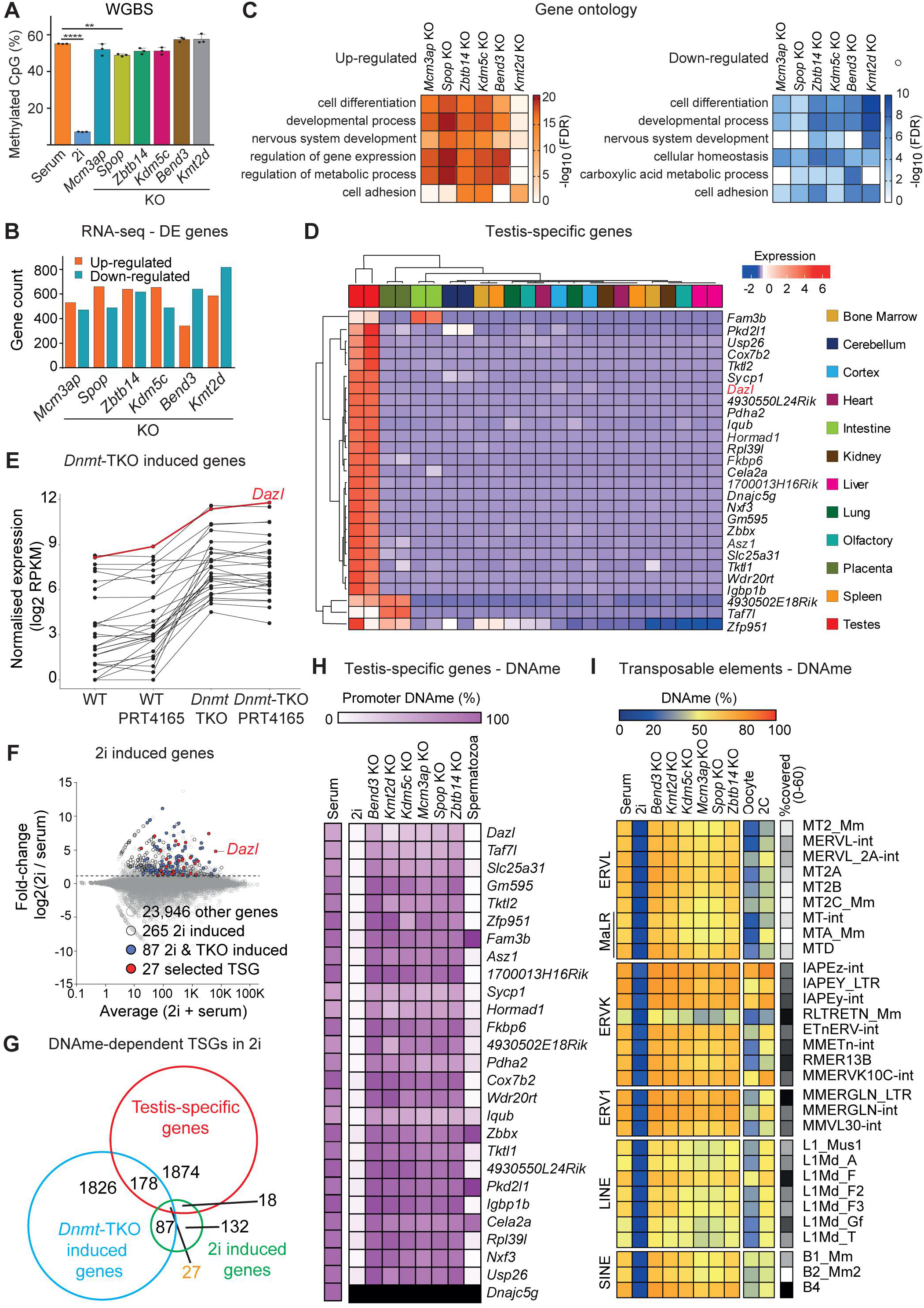
Regulation of DNA methylation-dependent testis-specific genes by novel epigenetic regulators. (A) Bar plot representing the average CpG methylations level measured by WGBS for 6 candidates KO and parental DASH mES cells cultured in Serum or 2i. (B) Bar plot representing the number of upregulated and downregulated genes for 6 candidates KO, compared to parental DASH mES cells. (C) Gene ontology (GO) terms (Slim Biological Process) significantly overrepresented among the upregulated (left) and downregulated (right) gene lists for 6 candidates KO, compared to parental DASH mES cells. (D) Hierarchically clustered heatmap depicting the elevated expression of 27 selected genes in testis compared to 11 other adult tissues (n=2 replicates per tissue). Transcriptome data was obtained from GSE29278 (Shen et al., 2012). (E) Dot plot showing the elevated expression of 27 DNA methylation-dependent testis-specific genes following the loss of repression by lack of DNA methylation in *Dnmt*-TKO complemented by limited induction under ncPRC1 inhibition by PRT4165. Transcriptome data was obtained from GSE101769 (Hill et al., 2018). (F) MA-plot showing the identification of 2i induced genes based on their increased expression under global hypomethylation in 2i culture conditions. (G) Venn diagram showing the overlap of three gene sets; testis-specific genes, *Dnmt*-KO induced genes, and 2i induced genes. 27 DNA methylation-dependent testis-specific genes were identified. (H) Heatmap depicting the average promoter DNA methylation (WGBS) of 27 selected testis-specific genes in 6 candidates KO (n=3, clones per KO) and parental DASH mES cells cultured in Serum or 2i. Spermatozoa methylation data was obtained from DDBJ DRA002402 (Kubo et al., 2015). (I) Heatmap depicting the average DNA methylation (WGBS) of SINE, LINE, ERV1, ERVK, and ERVL retrotransposons in 6 candidates KO (n=3, clones per KO) and parental DASH mES cells. The percent of individual elements covered by sufficient WGBS reads is indicated for each transposon family. Published data of oocyte and 2C embryos was obtained from GSE56697 (Wang et al., 2014).

**Figure S6:**
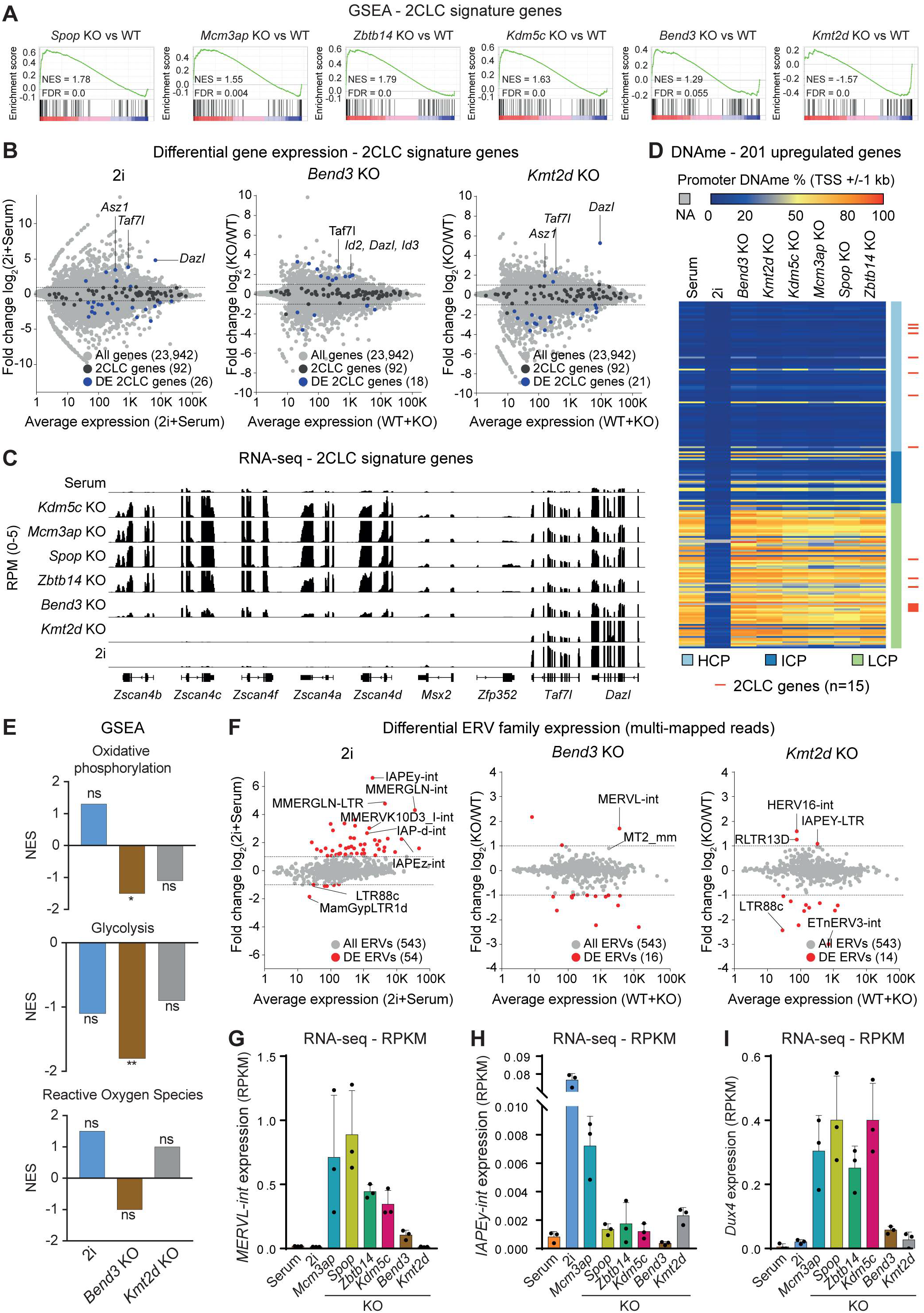
Knock-out of Spop, Mcm3ap, Kdm5c, and Zbtb14 induces Dux4, the master regulator of the 2CLC state. (A) GSEA plot showing significant upregulation of 2CLC markers in each of the *Spop* KO, *Mcm3ap* KO, *Kdm5c* KO, and *Zbtb14* KO, but not in *Bend3* KO and *Kmt2d* KO transcriptomic profiles. (B) MA-plots showing differentially expressed 2CLC genes in DASH mES cells cultured in 2i, *Bend3* KO, and *Kmt2d* KO. (C) Genome browser screenshots of expression profiles of selected 2CLC marker genes in 6 candidates KO and parental DASH mES cells cultured in Serum or 2i (tracks are an average of 3 replicates). (D) Heatmap depicting the promoter (TSS +/− 1kb) DNA methylation level of the 201 common upregulated genes classified by their promoter class, for 6 candidates KO and parental DASH mES cells cultured in Serum or 2i. (E) Bar plots showing Normalized Enrichment Score (NES) of selected metabolic pathways in DASH mES cells cultured in 2i, *Bend3* KO, and *Kmt2d* KO. *FDR<0.05, **FDR<0.01, ns: FDR > 0.05. (F) MA-plots showing differentially expressed ERV retrotransposons in DASH mES cells cultured in 2i, *Bend3* KO, and *Kmt2d* KO. (G-I) Normalized expression levels (RPKM) of *MERVL-int* (G), *IAPEy-int* (H), and *Dux4* (I) in 6 candidates KO (n=3, clones per KO) and parental DASH mES cells cultured in Serum or 2i.

**Figure S7:**
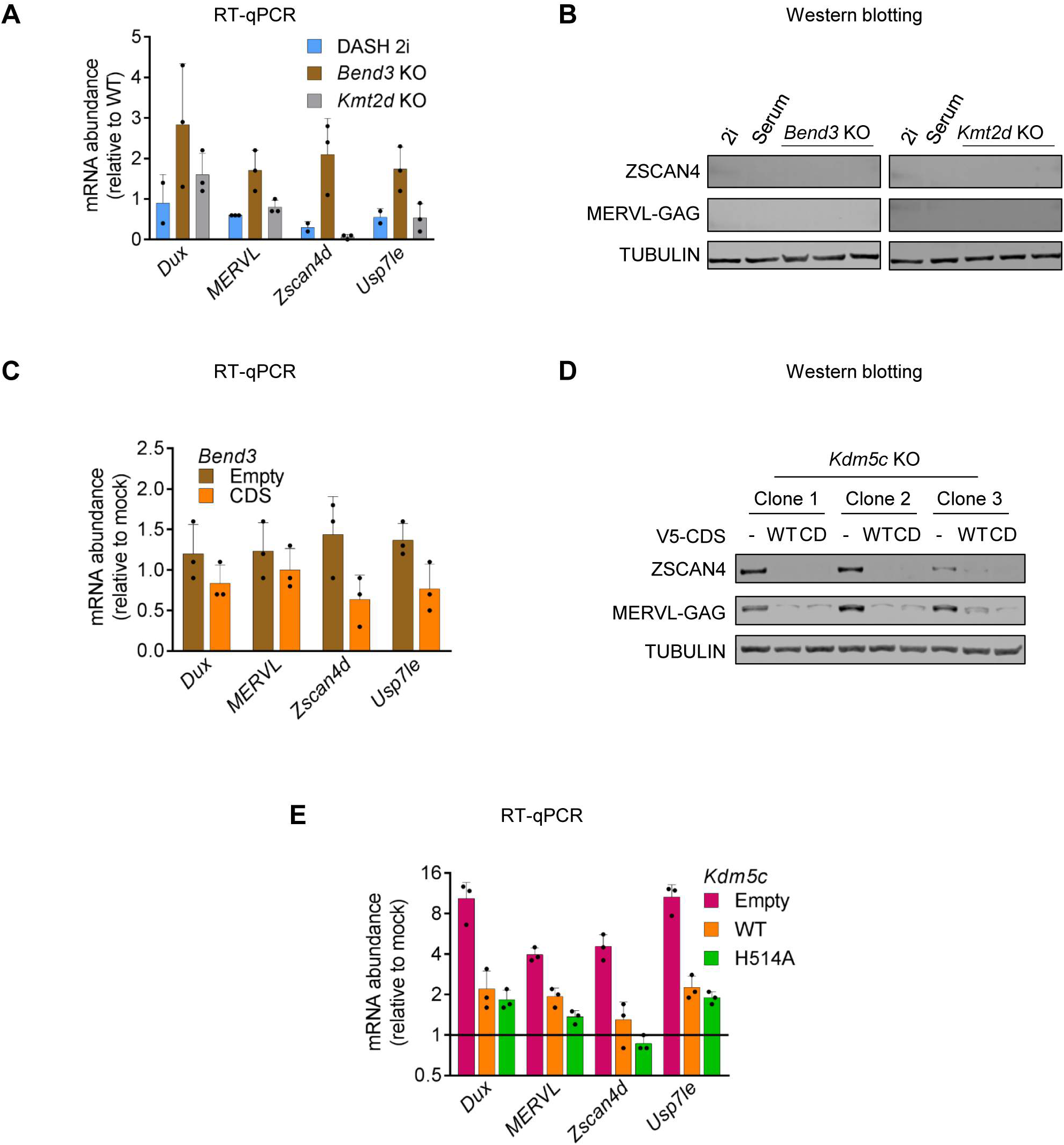
2CLC state is not induced in *Bend3* and *Kmt2d* KO. (A) RT-qPCR analysis of the expression of 2C-specific genes in DASH mES cells cultured in 2i, *Bend3* KO, and *Kmt2d* KO. (B) Western blot analysis of ZSCAN4 and MERVL-Gag for *Bend3* KO, *Kmt2d* KO, and parental DASH mES cells cultured in Serum or 2i. (C) RT-qPCR analysis of the expression of 2C-specific genes for *Bend3* KO (n=3, clones), transfected with an empty plasmid or their respective CDS, compared to the parental DASH mES cells. (D) Western blot analysis of ZSCAN4 and MERVL-Gag for *Kdm5c* KO clones, transfected with an empty plasmid, or a wild-type *Kdm5c* CDS, or a catalytically dead *Kdm5c* CDS (H514A). (E) RT-qPCR analysis of the expression of 2C-specific genes for *Kdm5c* KO clones, transfected with an empty plasmid, or a wild-type *Kdm5c* CDS, or a catalytically dead *Kdm5c* CDS (H514A).

## Accession number

The accession number for the deep sequencing data reported in this paper is GEO: GSE173573. Other deep sequencing data used are already published (see Methods).

## Author Contributions

NG and PAD conceived the project. NG, LY, and PAD planned the experiments. NG, and LY, performed experiments and analyzed the data. FM performed WGBS. LF performed LUMA. CD performed MeDIP. FB performed mass spectrometry. MD and BD performed Fluidigm. JRA performed RNA-seq and WGBS analysis. OK, ML, AS, KY, GC, and SB, performed bioinformatic analyses. NG, LY, and PAD wrote the manuscript. PAD, TI, and NG supervised the project. PAD acquired funding. All authors reviewed the manuscript.

## Competing interests statement

The authors declare no competing interests.

## Acknowledgments

We are very grateful to the following colleagues for useful advice: Allison Bardin, Claire Francastel, Claire Rougeulle, Sophie Polo, Pablo Navarro, Raphael Margueron, Petra Hajkova, Guillaume Velasco, Guillaume Filion, Miguel Casanova, Sainitin Donakonda, Michael Weber, Yoichi Shinkai, and Till Bartke. We thank the following colleagues for useful reagents: Déborah Bourc’his for J1 mESC, Nobuo Sakaguchi for a mouse GANP cDNA. We thank the Vectorology, Epigenetics and Microscopy platform at the Epigenetics and Cell Fate Unit, for providing access and technical advice. We acknowledge the ImagoSeine core facility of the Institut Jacques Monod, member of the France BioImaging (ANR-10-INBS-04) and the support of La ligue contre le Cancer (R03/75-79). Microfluidic RT-qPCR (Fluidigm) analysis was carried out on the qPCR-HD-Genomic Paris Centre Core Facility and was supported by grants from Région Ile-de-France, DIMBIO-RVT-INSERM-ADR-P11 n° 21016711. PAD is supported by Agence Nationale de la Recherche (PRCI INTEGER ANR-19-CE12-0030-01), LabEx “Who Am I?” (ANR-11-LABX-0071), Université de Paris IdEx (ANR-18-IDEX-0001) funded by the French Government through its “Investments for the Future” program, Fondation pour la Recherche Médicale, Fondation ARC (Programme Labellisé PGA1/RF20180206807). SB acknowledges support from Deutsche Forschungsgemeinschaft (DFG, 431163844). PAD, AS and GC were supported by grant RETROMET, ANR-16-CE12-0020, from Agence Nationale de la Recherche. JRA and MVCG were supported by Laboratoire d’excellence Who Am I? (Labex 11-LABX-0071) Emerging Teams Grant and by the European Research Council (ERC-StG-2019 DyNAmecs). This research was supported by Platform Project for Supporting Drug Discovery and Life Science Research (Basis for Supporting Innovative Drug Discovery and Life Science Research (BINDS)) from AMED under Grant Number JP20am0101103 (support number 2652).

## Materials and Methods

### Cell culture

J1 mouse embryonic stem cells (129S4/SvJae, XY) were cultured on gelatin-coated dishes in serum/LIF medium containing DMEM supplemented with 15% fetal bovine serum (FBS), non-essential amino acids (NEAA), penicillin/streptomycin, and 1000 U/ml leukemia inhibitory factor (LIF). When necessary, mouse embryonic stem cells (mESCs) were adapted to 2i/Vitamin C/LIF medium containing serum-free DMEM-F12 and Neurobasal media supplemented with 1% N2, 2% B27, 100 μg/ml ascorbic acid, 1 μM PD0325901, and 3 μM CHIR99021. Cells were incubated in a humidified atmosphere at 37°C under 5% CO2. For spontaneous differentiation, cells were seeded at clonal density in a serum medium without LIF.

### Cloning of sgRNA, transfection, and transduction in mES cells

The single-guide RNAs (sgRNAs) were designed using either Benchling/CRISPOR software or sequences were obtained from the Brie library (most enriched in the CRISPR KO screen). The gRNAs were cloned in either PX459 vector (Addgene #62988) or lentiCRISPRv2 (Addgene #52961), expressing both sgRNAs and Cas9 protein. Briefly, 10 µM sgRNA oligonucleotides were denatured at 95°C for 5 min, gradually cooled down to room temperature to anneal in a thermal cycler, and cloned into either BbsI-digested or Esp3I-digested plasmid by T4 DNA ligase (NEB) at 16°C overnight. The ligation mixture was then electroporated into NEB-Stbl3 bacteria; individual colonies were verified by sequencing. In some instances, the plasmids were transfected into DASH cells using an Amaxa 4D-Nucleofector (Lonza), according to the manufacturer’s instructions. The pmaxGFP plasmid was used as a positive control to estimate transfection efficiency. Alternatively, lentiviruses were generated and used for transduction. Production of lentiviral particles was performed by calcium-phosphate transfection of HEK293T with psPAX2 and pMD2.G plasmids, in a BSL3 tissue culture facility.

### Generation of DASH (<u>Da</u>zl-m<u>S</u>carlet-<u>H</u>ygromycin^R^) reporter cell line

The reporter cassette comprising of P2A-*mScarlet*-T2A-*Hygro^R^* (synthesized by GenScript) was inserted in-frame within the exon 6 of mouse *Dazl* gene. The reporter cassette encoded proteins useful for phenotypic assays and cell selection: the bright red-fluorescent protein, mScarlet, and the hygromycin-resistance Hygro^R^. The two proteins expressed by the cassette were separated from each other, and the N-terminus of DAZL, by self-cleaving 2A peptides (P2A, T2A). The reporter cassette also included linker regions between all functional proteins and a stop codon at the end, thus disrupting the inserted endogenous *Dazl* allele. The exon 6 was chosen as the insertion site since it met several criteria, such as being a frame-shifting exon present in all mRNA isoforms. The cassette was flanked by *Dazl* homology arms (HA) corresponding to endogenous intron 5-exon 6 and intron 6 sequences respectively. Protospacer Adjacent Motif (PAM) sites of the two sgRNAs targeting *Dazl* exon 6 were mutated in the homology arms to prevent re-cutting of Cas9 after the insertion of the cassette through homologous recombination. The synthesized cassette was cloned into pUC57-Simple. The two sgRNAs targeting *Dazl* exon 6 were cloned into the pSpCas9(BB)-2A-GFP backbone (Addgene #48138). The knock-in led to the generation of a reporter cell line named *Dazl*-mScarlet-Hygromycin^R^ (DASH), carrying a homologous integration of the reporter cassette in one of the alleles of *Dazl*, confirmed by DNA sequencing.

### Uhrf1 KO, validation of DASH mES cells

To test the utility of the DASH cell line, two sgRNAs targeting *Uhrf1-*exon2 were designed, cloned in the PX459 vector, transfected into DASH cells to test the reporter, followed by a short pulse of 1 µg/ml puromycin for 24 h, and knockout efficiency was assessed by western blotting.

### CRISPR KO screen: amplification of sgRNA library, lentiviral transduction, and sample collection

We performed a genome-wide CRISPR knock-out (KO) screen using the lentiviral Brie sgRNA library comprising 4 sgRNAs per protein-coding gene (Addgene #73632) (Doench et al., 2016). The sgRNA library was amplified and sequenced confirming an equal representation of the ∼80,000 sgRNAs, as expected and lentiviruses were produced as described above. The screen in DASH mESCs was performed in two biological replicates, cells were transduced with a lentiviral plasmid pool at a multiplicity of infection (MOI) ∼0.1. After 48 h, transduced cells were positively selected with 2 µg/ml puromycin for 5 days. Coverage was 150X (150 transduced cells/gRNA) for each biological replicate. Sequencing of the Puro^R^ sample confirmed a comprehensive representation of the sgRNA library in the screen.. Following this, cells were initially selected with a non-lethal dose of 50 µg/ml hygromycin for 3 days followed by additional selection for 11 days at a lethal dose of hygromycin (125 µg/ml). Cell medium was changed every day with fresh antibiotics. Three weeks post-infection, hygromycin-resistant cells were sorted by FACS for mScarlet expression. Samples were collected at multiple time points to allow comparisons with the end-point sample, mScarlet+ cells, to track sgRNA diversity and their enrichment. Puromycin-resistant (Puro^R^) cells represented the initial sgRNA diversity and depth among the transduced cells. Two samples were collected during hygromycin selection: first, at a starting non-lethal dose (Hygro^50^), and then at a lethal dose (Hygro^R^). Finally, mScarlet expressing cells were collected by FACS sorting. For sample collection, at each mentioned time point, all cells in different dishes were pooled and about 25% of the total cells were harvested, to avoid creating a bottleneck to the representation of sgRNA diversity.

### CRISPR KO screen: sequencing and analysis

Genomic DNA was isolated from cells using AllPrep DNA/RNA Mini Kit (Qiagen) following the recommended protocol. For the sgRNA plasmid library, 50 ng DNA was used for each reaction whereas 200 ng of genomic DNA was used per reaction. Briefly, PCR was performed with Platinum Taq polymerase (Thermo Fisher Scientific), employing a pool of P5 primers (designed in the lab) and a unique P7 barcode primer (primer sequences are listed in Supplementary File 2). The PCR conditions were: initial denaturation at 94°C for 4 min; 28 cycles of denaturation at 94°C for 30 s, annealing at 53°C for 30 s and extension at 72°C for 30 s per kb; final extension at 72°C for 10 min. For each sample, four 50 µl PCR reactions were performed and pooled. The PCR products were retrieved using QIAquick PCR Purification Kit (Qiagen) and verified for the right amplicon on an agarose gel. The DNA was further purified using AMPure XP (Beckman Coulter). Libraries were sequenced on an Illumina HiSeq 1500 in single-end (SE) 100 bp output mode. The sgRNA distribution and enrichment at different time points were analyzed by MAGeCK workflow (Chen et al., 2018). A statistical threshold of p-value < 0.05% resulted in a list of 40 candidates whose knockout led to the expression of mScarlet and Hygro^R^ in DASH mES cells. The list of enriched genes is available in Supplementary File 3.

### Generation of individual gene KO and clonal cell lines

To validate 24 statistically significant candidates, individual KO in DASH mESCs were generated using the top two most efficient sgRNAs (as determined by MAGeCK analysis, Supplementary File 3). During lentiviral production, both sgRNA plasmids targeting the same gene were mixed to increase knockout efficiency. Transduced cells were selected with 2 µg/ml puromycin for 3 days; followed by hygromycin selection (50 µg/ml for 3 days, and 125 µg/ml for the next 7 days). Further, for the chosen 6 candidates, i.e., *Bend3, Kdm5c, Kmt2d, Mcm3ap, Spop,* and *Zbtb14*, three independent clonal KO lines were established.

### Rescue experiments, piggyBac system

For rescue experiments, the coding sequence (CDS) of candidates was synthesized (for *Bend3, Spop,* and *Zbtb14)* (GenScript), amplified from cDNA (*Kdm5c*), or obtained from colleagues (for *Mcm3ap*, kindly shared by Prof. N. Sakaguchi, Singh et al., 2013). In all cases, silent mutations were incorporated either within the PAM and/or sgRNA sequence to block targeting by the active cognate sgRNAs in the KO clonal cell lines. For *Kdm5c*, an additional catalytically-dead (CD) mutant plasmid with the H514A mutation was generated. These CDS sequences were cloned into a piggyBac vector and co-transfected with PB transposase for stable insertion (Fukuda et al., 2018). Empty piggyBac vector served as a control. Transfected cells were selected with 5 µg/ml blasticidin for 5 days, and processed for phenotypic and molecular assays.

### Digital droplet PCR (ddPCR)

The PCR reaction mixture composed of 2X EvaGreen ddPCR Supermix (Bio-Rad), primers at a final concentration of 100 nM and 10 ng of template DNA were partitioned into up to 20,000 droplets by water-oil emulsion. After droplet generation, a regular PCR was performed with the following conditions: 95°C for 5 min; 95°C for 30 s and 60°C for 1 min (40 cycles); 4°C for 5 min, 90°C for 5 min, 4°C hold. For all steps, a ramp rate of 2°C/s was used. Cycled droplets were read individually (Bio-Rad QX-200 droplet reader). Each run included technical duplicates and no-template controls. Primer sequences are listed in Supplementary File 2.

### RT-qPCR

Total RNA was extracted from cells with RNeasy Plus Mini kit (Qiagen) according to the manufacturer’s instructions and quantified using Qubit RNA BR Assay kit on Qubit 2.0 Fluorometer (Thermo Fisher Scientific). One microgram of total RNA was reverse transcribed using SuperScript IV Reverse Transcriptase (Thermo Fisher Scientific) and Oligo dT primers (Promega). RT-qPCR was performed using Power SYBR Green (Applied Biosystems) on a ViiA 7 Real-Time PCR System (Thermo Fisher Scientific). *Actinb*, *Ppia,* and *Rplp0* were used for normalization. Primer sequences are available in Supplementary File 2.

### Microfluidic RT-qPCR (Fluidigm): cDNA synthesis, RT-qPCR, and analysis

Total RNA was extracted and quantified as described above. RNA quality and integrity were analyzed by capillary electrophoresis with the Fragment Analyzer (Agilent Technologies) to calculate the RNA quality number (RQN) for each sample. Defined on a scale ranging from 1 to 10, the mean RQN of the 52 samples was 9.9, indicating very good RNA quality. The cDNA synthesis was performed using Reverse Transcription Master Mix (Fluidigm) according to the manufacturer’s protocol in a final volume of 5 μl containing 200 ng total RNA. Reverse transcription was performed using a Nexus Thermocycler (Eppendorf) with the following parameters: 5 min at 25°C, 30 min at 42°C followed by heat-inactivation of reverse transcriptase for 5 min at 85°C and immediately cooled to 4°C. cDNA samples were diluted 5X by adding 20 μl of low TE buffer (10 mM Tris, 0.1 mM EDTA, pH 8.0) (TEKNOVA) and stored at −20°C before specific target pre-amplification. Diluted cDNA was used for multiplex pre-amplification in a total volume of 5 μl containing 1 µl of 5X PreAmp Master Mix (Fluidigm), 1.25 µl of cDNA, 1.25 µl of pooled assay (Exiqon) with an original concentration of each assay of 10 µM and 1.5 μl of nuclease-free water. The cDNA samples were subjected to pre-amplification with the following conditions: 95°C for 2 min, followed by 10 cycles at 95°C for 15 s and 60°C for 4 min. The pre-amplified cDNA was diluted 5X by adding 20 µl of low TE buffer and stored at - 20°C before RT-qPCR. The expression of 48 target genes was quantified in 52 samples by quantitative PCR using the high-throughput platform BioMark HD System on two 48×48 GE Dynamic Arrays (Fluidigm). Pre-amplified cDNA was used in a total volume of 15 μl containing 7.5 μl of 2X Gene Expression PCR Master Mix (Thermo Fisher Scientific), 3 μl of pre-amplified cDNA, 0.75 μl of 20X EvaGreen (Biotium), 1.5 μl of each primer (Exiqon) and 2.25 µl nuclease-free water. Thermal conditions for qPCR were: 50°C for 2 min before initial heat-activation of DNA polymerase at 95°C for 10 min, followed by 40 cycles at 95°C for 15 s and 60°C for 1 min. Melt curve analysis was performed with an increase of 0.2°C/s, starting from 65°C for 5 s and ending at 95°C. The parameters of the thermocycler were set with ROX as a passive reference and a single probe EvaGreen as a fluorescent detector. To determine the quantification cycle Cq, data were processed by an automatic threshold for each assay, with linear derivative baseline correction using BioMark Real-Time PCR Analysis Software 4.0.1 (Fluidigm). The quality threshold was set at the default setting of 0.65. Second data analysis was performed using R software to measure the relative gene expression using the comparative Cq method with the efficiency corrected method of Pfaffl, after normalization with reference genes (Cq mean of *Actinb*, *Ppia,* and *Rplp0*). Fold change between experimental and control groups was calculated for each sample as the difference of Cq between reference genes and the gene of interest (GOI) in control and experimental conditions.

### Western blotting

Cells were harvested and lysed in RIPA buffer (Sigma) with protease inhibitor cocktail (Thermo Fisher Scientific), sonicated with a series of 30s ON / 30s OFF for 5 min on a Bioruptor (Diagenode), and centrifuged at 16,000 g for 5 min at 4°C. The supernatant was collected and quantified by BCA assay (Thermo Fisher Scientific). Thirty microgram protein extract per sample was mixed with NuPage 4X LDS Sample Buffer and 10X Sample Reducing Agent (Thermo Fisher Scientific) and denatured at 95°C for 5 min. Samples were resolved on a pre-cast SDS-PAGE 4-12% gradient gel (Thermo Fisher Scientific) with 120V electrophoresis for 90 min and blotted onto a nitrocellulose membrane (Millipore). The membrane was blocked with 5% fat-free milk/PBS at RT for 1 h, then incubated overnight at 4°C with appropriate primary antibodies. After three washes with PBS/0.1% Tween20, the membranes were incubated with the cognate fluorescent secondary antibodies and revealed in the LI-COR Odyssey-Fc imaging system. The following antibodies were used in this study: α-DAZL (Abcam #ab34139, 1:500), α-DNMT3A (Cell Signaling #3598, 1:500), α-DNMT3B (Novus #56514, 1:500), α-UHRF1 (SCBT #98817; 1:1000), α-V5 (Abcam #ab206566, 1:1000), α-MuERVL-Gag (HuaBio #ER50102, 1:1000), α-ZSCAN4 (Merck #AB4340, 1:5000), α-TUBULIN (Abcam #7291; 1:10000), α-GAPDH (Abcam #ab9485, 1:10000). The following secondary antibodies were used in this study: IRDye 800CW Donkey α-Rabbit (Licor #926-32213, 1:5000) and IRDye 680RD Donkey α-Mouse (Licor #926-68072, 1:5000).

### Isolation of genomic DNA

Genomic DNA was isolated from cells using overnight 200 μg/ml proteinase K treatment at 55°C followed by 20 μg/ml RNase A treatment at 37°C for 1 h and extracted by standard phenol/chloroform/alcohol method. Alternatively, genomic DNA was isolated from cells using QIAamp DNA Mini kit (Qiagen), following the manufacturer’s instructions. Genomic DNA was resuspended in water and quantified with Qubit dsDNA BR Assay kit on Qubit 2.0 Fluorometer (Thermo Fisher Scientific). DNA integrity was assessed with Genomic DNA ScreenTape on TapeStation system (Agilent) and samples with DNA Integrity Number > 9 were used for subsequent analysis.

### DNA methylation analysis: Methylated DNA Immunoprecipitation (MeDIP)

MeDIP was performed using the Auto MeDIP Kit on an automated platform SX-8G IP–Star Compact (Diagenode). Briefly, 2.5 μg of DNA was sheared using a Bioruptor Pico to approximately 500-bp fragments, as assessed with D5000 ScreenTape (Agilent). Cycle conditions were as follows: 15 s ON / 90 s OFF, repeated 6 times. A portion of sheared DNA (10%) was kept as input and the rest of the sheared DNA was immunoprecipitated with α-5-methylcytosine antibody (Diagenode), bound to magnetic beads, and was isolated. qPCR for selected genomic loci was performed and efficiency was calculated as % (me-DNA-IP/total input). Primer sequences are listed in Supplementary File 2.

### DNA methylation analysis: Luminometric methylation assay (LUMA)

To assess global CpG methylation, 500 ng of genomic DNA was digested with MspI+EcoRI and HpaII+EcoRI (NEB) in parallel reactions, EcoRI was included as an internal reference. CpG methylation percentage is defined as the HpaII/MspI ratio. Samples were analyzed using PyroMark Q24 Advanced pyrosequencer

### DNA methylation analysis: LC-MS/MS

The genomic DNA was extracted as described above with an additional step of digestion with RNase A. One microgram of DNA was treated with 10U DNA Degradase Plus (ZymoResearch) at 37°C for 4 h. After enzyme inactivation at 70°C for 20 min, the solution was filtered with Amicon Ultra-0.5 mL 10 K centrifugal filters (Merck Millipore). The reaction mix retained in the centrifugal filter was processed for LC-MS/MS analysis. Analysis of global levels of 5-mdC and 5-hmdC were performed on a Q exactive mass spectrometer (Thermo Fisher Scientific). It was equipped with an electrospray ionization source (H-ESI II Probe) coupled with an Ultimate 3000 RS HPLC (Thermo Fisher Scientific). Digested DNA was injected onto a ThermoFisher Hypersil Gold aQ chromatography column (100 mm * 2.1 mm, 1.9 µm particle size) heated at 30°C. The flow rate was set at 0.3 ml/min and run with an isocratic eluent of 1% acetonitrile in water with 0.1% formic acid for 10 minutes. Parent ions were fragmented in positive ion mode with 10% normalized collision energy in parallel-reaction monitoring (PRM) mode. MS2 resolution was 17,500 with an AGC target of 2e5, a maximum injection time of 50 ms, and an isolation window of 1.0 m/z. The inclusion list contained the following masses: dC (228.1), 5-mdC (242.1) and 5-hmdC (258.1). Extracted ion chromatograms of base fragments (±5ppm) were used for detection and quantification (112.0506 Da for dC; 126.0662 Da for 5-mdC; 142.0609 for 5-hmdC). Calibration curves were previously generated using synthetic standards in the ranges of 0.2 to 10 pmol injected for dC and 0.02 to 10 pmol for 5mdC and 5hmdC. Results are expressed as a % of total dC.

### Flow cytometry

The number of cells expressing mScarlet was determined by flow cytometry using a yellow laser (561 nm) of the Influx or FACSAria Fusion cell sorter (BD Biosciences) at the ImagoSeine core facility (Institut Jacques Monod). A threshold on mScarlet signal intensity (subtracting background fluorescence from non-reporter wild-type mESC) was used to determine the proportion of positive cells. Data were analyzed with FlowJo software.

### Crystal violet staining

Cells were seeded at the same density in all wells and grown with or without hygromycin for 7 days. Surviving cells were fixed with absolute ethanol for 15 min, stained with 1% Crystal violet dye (Sigma) for 30 min, and washed extensively with water to remove the unbound stain.

### Gene ontology analysis

Gene ontology (GO) analysis was performed using UniprotKB ID in the DAVID v6.8 tool for the 40 significant hits from the genome-wide CRISPR KO screen. FDR values for all significantly overrepresented GO terms from BP_direct were displayed. GO analysis was performed using the PANTHER v16.0 tool for differentially expressed genes identified by RNA-seq in this study. FDR values for selected significantly overrepresented GO terms from Slim Biological Process were displayed. Complete GO analysis is available in Supplementary File 4.

### Protein-Protein Interaction Network analysis

Protein-protein interactions for the 40 significant hits from the genome-wide CRISPR KO screen were performed using the STRING v11.0 tool. Interactions were computed using default parameters and network edges with a confidence score > 0.6 were displayed. Network visualization was edited with Cytoscape software.

### Identification of CRISPR-induced mutation

Genomic DNA was isolated as described above. Amplicons spanning sgRNA-targeted regions were generated by PCR, using 100 ng of genomic DNA, 100 nM primers (listed in Supplementary File 2), and Platinum Taq polymerase (Thermo Fisher Scientific) in 50 μl reactions. The conditions were: initial denaturation at 94°C for 4 min; 35 cycles of denaturation at 94°C for 30 s, annealing at 58°C for 30 s and extension at 72°C for 30s per kb; final extension at 72°C for 10 min. Amplicons ranging from 291 to 481 bp were generated for deep-sequencing; the library for sequencing was generated using NEBNext Ultra II DNA Library Prep Kit for Illumina (NEB) following the manufacturer’s recommendations. Libraries were sequenced by Novogene on NovaSeq 6000 generating 250 bp paired-end reads. The filtered fastq files were used as input file for AmpliCan software in R. To avoid other library contamination, AmpliCan was run with “fastqfiles = 0” and “primer_mismatch = 0” setting. The other parameters for AmpliCan were left as default. The identified mutations are listed in Supplementary File 1.

### RNA-sequencing: Library preparation for transcriptome sequencing

A total amount of 1 μg total RNA per sample was used as input material for the RNA sample preparations. RNA samples were spiked with ERCC RNA Spike-In Mix (Thermo Fisher Scientific). Sequencing libraries were generated using NEBNext UltraTM RNA Library Prep Kit for Illumina (NEB) following the manufacturer’s recommendations. Briefly, mRNA was purified from total RNA using poly-T oligo-attached magnetic beads. Fragmentation was carried out using divalent cations under elevated temperature in NEBNext First Strand Synthesis Reaction Buffer (5X). First-strand cDNA was synthesized using a random hexamer primer and M-MuLV Reverse Transcriptase (RNase H-). Second strand cDNA synthesis was subsequently performed using DNA Polymerase I and RNase H. In the reaction buffer, dNTPs with dTTP were replaced by dUTP. The remaining overhangs were converted into blunt ends via exonuclease/polymerase activities. After adenylation of 3’ ends of DNA fragments, NEBNext Adaptor with hairpin loop structure was ligated to prepare for hybridization. To select cDNA fragments of preferentially 250-300 bp in length, the library fragments were purified with the AMPure XP system (Beckman Coulter). Then 3 μl USER Enzyme (NEB) was used with size-selected, adaptor-ligated cDNA at 37°C for 15 min followed by 5 min at 95°C before PCR. Then PCR was performed with Phusion High-Fidelity DNA polymerase, Universal PCR primers, and Index (X) Primer. At last, products were purified (AMPure XP system) and library quality was assessed on the Agilent Bioanalyzer 2100 system.

### RNA-sequencing: read alignment

FASTQ reads were trimmed using Trimmomatic (v0.39) and parameters: ILLUMINACLIP:adapters.fa:2:30:10 SLIDINGWINDOW:4:20 MINLEN:36. Read pairs that survived trimming were aligned to the mouse reference genome (build mm10) using STAR (v2.7.5c) and default single-pass parameters. PCR duplicate read alignments were flagged using Picard-tools (2019) MarkDuplicates (v2.23.4). Uniquely aligned, non-PCR-duplicate reads were kept for downstream analysis using Samtools view (v1.10) and parameters: -q 255 -F 1540. Gene expression values were calculated over the mm10 NCBI RefSeq Genes annotation using VisRseq (v0.9.12) and normalized per million aligned reads per transcript length in kilobases (RPKM). Bigwig files were generated using deeptools bamCoverage (v3.3.0) using counts per million (CPM) normalization and visualized in IGV (v2.8.9).

### RNA-seq: Differential expression, PCA plots, and heatmaps

DESeq2 (v1.30.0) was employed using apeglm LFC shrinkage to calculate differential expression. Genes or transposable elements were categorized as significantly differentially expressed if they showed an expression fold-change >=2 and associated adjusted p-value <0.01. PCA plots were generated using variance-stabilized RNA-seq read counts using ‘varianceStabilizingTransformation’, and plotted using the DESeq2 function ‘plotPCA’ and visualized using the R package ‘ggplot2’ (v3.3.3) in R (v4.0). Heatmaps were generated using Morpheus (https://software.broadinstitute.org/morpheus).

### Gene set enrichment analysis (GSEA)

Gene set enrichment analysis was performed using GSEA (v4.1.0) and default parameters (1000 permutations, permutation type = gene_set. Selected significant terms from Hallmark gene sets (n=50 “h.all.v7.4.symbols.gmt”) were displayed. Complete GSEA analysis is available in Supplementary File 5.

### RNA-seq: Transposable element quantification

RepeatMasker (last updated 2012-02-06) was downloaded from the UCSC Table Browser. To measure the expression of transposable element families, PCR duplicates were removed and all reads, including uniquely mapped and multi-mapped reads, were enumerated using VisRseq. Multi-mapped reads were counted once, and all individual elements were aggregated to calculate family-wide expression in read count (for differential expression analysis) and RPKM values (for heatmap illustrations, available in Supplementary File 6).

### RNA-seq: MERVL/MT2_Mm analysis

MERVL internal sequences and their MT2_Mm LTR promoters were extracted from the RepeatMasker annotation (last updated 2012-02-06). Internal sequences and their LTRs located within 88 bp were merged into a single element using bedtools merge (v2.27.0) to account for an 87 bp insertion of a related ORR1A3 element. Elements were categorized as full-length MERVL elements if they contained both LTR elements and internal sequences and spanned >6000bp. MT2_Mm elements under 500 bp in length were defined as “Solo-LTRs”. All other elements, such as those composed of MERVL internal sequences and only one LTR, were categorized as “other”. Genome-wide mappability scores were calculated using iGEM (v1.315) and parameters: K_MER_SIZE=300 MAX_MISMATCHES=0.04 and the mappability of each MERVL element was calculated using VisRseq. A list of MERVL elements that generate chimeric transcripts was downloaded from (Macfarlan et al., 2012) and mapped onto the mm10 genome using UCSC LiftOver. To measure individual transposable element expression, only uniquely aligned, non-PCR duplicate reads were counted. Elements were grouped and sorted by K-medoid clustering on log10-transformed RPKM values using the R package “cluster” and VisRseq.

### RNA-seq: Dux expression analysis

Due to the contentious nature of the mouse Dux locus in the mm10 build, we aligned RNA-seq reads directly to the Dux repeat (AM398147.1), as described previously (Eckersley-Maslin et al., 2019). Duplicate alignments were ignored and values were reported as reads per million sequenced (RPM).

### Whole-genome-bisulfite sequencing (WGBS)

Genomic DNA was extracted as described for LC-MS/MS. The library preparation for WGBS was performed with the tPBAT protocol described previously (Miura et al., 2012, Miura et al., 2019). One hundred nanograms of genomic DNA spiked with 1% (w/w) of unmethylated lambda DNA (Promega) was used for the library preparation. The sequencing was performed by Macrogen Japan Inc. using the HiSeq X Ten system. We assigned 8 lanes for the analysis of 20 samples. The sequenced reads were mapped with BMap and summarized with an in-house pipeline as described previously (Miura et al., 2019), with custom scripts archived using GitHub (https://github.com/FumihitoMiura/Project-2). The basic metrics of the methylome data are provided in Supplementary File 7. DNA methylation levels over CpGs covered by at least 5 sequencing reads were averaged over the following regions of interest: genome-wide 2kb bins, CpG islands (cpgIslandExt, n=16,023, last updated: 2012-02-09), enhancer elements (Ensembl Regulatory features release 81, n=73,796), promoters (NCBI Refseq, TSS +/− 1kb, n=24,371), gene bodies and transposable elements (RepeatMaster, n=5,147,736). Promoters (TSS +/− 1kb) were classified by CpG density as described previously (Weber et al., 2007).

### Identification of Testis-specific genes

Testis-specific genes were identified using public mouse tissue expression data (GSE29278) (Shen et al., 2012). Median expression was computed for all genes, in testis and in 11 other adult tissues. Genes significantly more expressed in testis compared to other tissues were identified with the following thresholds: log_2_(FC) > 2 and FDR < 0.1. Almost all of them (97%) had a median expression in testis greater than the average median expression in all other tissues: these were defined as testis-specific genes (n = 2687, Supplementary File 8).

### Defining Dnmt-TKO upregulated genes

*Dnmt*-TKO upregulated genes were identified using public RNA-seq data of *Dnmt*-TKO mESC (GSE101769) (Hill et al., 2018). Genes significantly induced upon *Dnmt*-TKO (versus the corresponding WT strain) were identified with the following thresholds: log_2_(FC) > 1 & FDR < 0.01. The contribution of DNA methylation to gene silencing was also taken into account in the absence of PRC1, by comparing *Dnmt*-TKO versus WT, both under PRT4165 treatment (log_2_(FC) > 1 & FDR < 0.01). The overlap of these two genelists, except for genes displaying an additive regulation by both pathways (n = 54), corresponded to *Dnmt*-TKO upregulated genes (n = 2390, Supplementary File 8).

### Defining 2i-induced genes

Promoter DNA methylation and gene expression (using RNA-seq and WGBS data from this study) were taken into account to define 2i-induced genes. Genes with promoters that displayed over 35% DNA methylation in serum-grown ESCs and less than 10% DNA methylation in 2i-grown cells were identified. Secondly, genes that showed over 2-fold upregulation in 2i-grown cells versus serum-grown cells were identified using DESeq2 (log_2_(FC) > 1 & FDR < 0.01). 2i-induced genes were defined as the intersection of both lists (available in Supplementary File 8).

### Identification of epigenetically regulated testis-specific genes

Intervene app was used to plot the overlap between 2i-induced genes, testis-specific genes and *Dnmt*-TKO upregulated genes (listed in Supplementary File 8), corresponding to 27 testis-specific genes, including *Dazl*, sensitive to epigenetic regulation.

## Notes

### Competing Interest Statement

The authors have declared no competing interest.

